# Chronic Pb^2+^ Exposure Causes Hippocampal Network Hypersynchrony, Absence Seizures, and Sensorimotor Deficits

**DOI:** 10.1101/2020.06.30.181149

**Authors:** Nathan W. Schultheiss, Jennifer L. McGlothan, Tomás R. Guilarte, Timothy A. Allen

**Affiliations:** Department of Psychology, Florida International University, Miami, FL, 33199; Department of Environmental Health Sciences, Robert Stempel College of Public Health, Florida International University, Miami, FL, 33199; Brain, Behavior & the Environment Program, Florida International University, Miami, FL, 33199; Cognitive Neuroscience Program, Florida International University, Miami, FL, 33199; Brain Institute, Nicklaus Children’s Hospital, Miami, FL 33135

**Keywords:** hippocampus, gamma, schizophrenia, lead neurotoxicity, pre-pulse inhibition

## Abstract

Chronic early-life lead (Pb^2+^) exposure contributes to an array of cognitive and behavioral dysfunctions, including impaired attention, memory, and intellectual abilities, in addition to increased social delinquency. Notably, Pb^2+^ exposure is an environmental risk factor for adult psychopathologies, including schizophrenia and epilepsy. Neurobiologically, Pb^2+^ is a potent N-methyl-D-aspartate receptor (NMDAR) antagonist, and exposure during early life elicits a cascade of cellular neurotoxic effects that alter neurodevelopmental trajectories. This includes reduced parvalbumin-expressing interneurons in the hippocampus (HC) and altered synaptic transmission. Little is known, however, about the impact of chronic Pb^2+^ exposure on HC network dynamics, which link cellular-molecular effects with cognitive-behavioral consequences. Here, we tested the impact of chronic Pb^2+^ exposure on the HC local field potential (LFP) in freely behaving rats. We found that Pb^2+^ exposure (1) caused a striking level of theta rhythmic hypersynchrony, (2) amplified fast gamma synchronization, (3) disrupted behavioral modifications of theta and fast gamma, and (4) exacerbated absence seizures appearing in the LFP as spike-wave discharges (SWDs) at theta frequencies. Each of these rhythmic changes in the HC network was related to exploratory movements in the open field. HC network alterations like these have also been linked to impaired prepulse inhibition of the acoustic startle reflex (PPI). Thus, next, we tested the effect of Pb^2+^ exposure on PPI. We found that adult males (PN50 and 120), but neither females nor juvenile males, showed reduced PPI, recapitulating sex dependencies on PPI disruptions in schizophrenics. Altogether, these results are consistent with the hypothesis that chronic early-life Pb^2+^ exposure causes dysfunction in the rhythmic network coordination of the HC, limiting processing, and helping to account for cognitive deficits.

## Introduction

Lead (Pb^2+^) is a pervasive pollutant and potentially lethal toxin (Pb^2+^ tox profile for ATSDR, 2020) that accounts for 0.6% of the total global cost of disease (Sample, 2021). Even low-level exposure to Pb^2+^ during early childhood causes permanent neurological dysfunctions and may be a risk factor for schizophrenia (SZ)(Guilarte et al., 2012) or epilepsy (Sasmaz et al., 2003). Since the late 1970s, public health policy to eliminate Pb^2+^ from gasoline, paint, and other consumer products has helped lower blood Pb^2+^ levels (BLL) in the general US population (Dignam et al., 2019). Still, a shocking proportion of children and adults, particularly in disadvantaged urban communities, continue to be exposed to Pb^2+^ due to the legacy of millions of tons of environmental contamination (Bernard & McGeehin, 2003; Yeter et al., 2020). As recently as 2002, the healthcare cost-burden of Pb^2+^ exposure in the US was 2.2% of the total national cost of healthcare (Wakefield, 2002); but numerous *ongoing* cases of significant Pb^2+^ exposure from municipally-regulated resources, e.g. public water (Hanna-Attisha et al., 2016) and affordable housing (Garnett & Iannuzzi, 2020), show that exposure risks are more extensive than previously appreciated (https://www.reuters.com/investigates/special-report/usa-lead-testing/). Recent US estimates of children with BLLs above the current Centers for Disease Control (CDC) reference level (10 μg/dL) exceed one-quarter million (e.g., Egan et al., 2021). Global estimates exceed one-half billion children, as detailed in the 2020 UNICEF report entitled: *The Toxic Truth-Children’s Exposure to Lead Pollution Undermines a Generation of Future Potential* (UNICEF, 2020).

Childhood exposure to Pb^2+^ alters brain development, leading to impoverished cognitive abilities, as well as social, emotional, and behavioral dysregulations that persist in adulthood (Lanphear et al., 2005; 2006; Canfield et al., 2003; Needleman et al., 2002; 2004; Miranda et al., 2007; Caito & Aschner, 2017). Young children are disproportionately vulnerable to Pb^2+^ neurotoxicity (Canfield et al., 2003) because their predisposition for exploratory behaviors increases exposures, and the immature blood-brain barrier is permissive of Pb^2+^ penetration, leading to markedly higher brain concentrations (Barltrop, 1969; Rossouw et al., 1987). Chelation therapy to reduce elevated BLLs can help mitigate the Pb^2+^ burden on the body, but it does not affect cognitive or neuropsychological impairments (Rogan et al., 2001, Dietrich et al., 2004).

Notably, there are no proven therapies to ameliorate the neurotoxic damage caused by childhood Pb^2+^ exposure once it has been sustained, and exposure prevention is the only available public health strategy (CDC, 2021).

In the brain, Pb^2+^ is a potent antagonist of NMDARs at excitatory synapses in the HC and medial prefrontal cortex (mPFC) (Guilarte & Miceli, 1993; Guilarte, 1997; 2023). Pb^2+^ binds to allosteric sites on NMDARs (Gavazzo et al., 2008), causing receptor hypoactivity and reduced calcium influx (Hashemzadeh- Gargari & Guilarte, 1999), diminishing kinase activity and nuclear CREB phosphorylation (Nihei et al., 2000; Toscano et al., 2002; Toscano et al., 2005; Toscano & Guilarte, 2005); and attenuating synaptic levels of brain-derived neurotrophic factor (BDNF) (Neal et al., 2010; Neal & Guilarte, 2010; Stansfield et al., 2012). Significantly, developmental Pb^2+^ exposure alters the receptor-subunit composition of NMDARs (Guilarte & McGlothan, 1998; Nihei et al., 2000), leading to impaired long-term potentiation (LTP) (Nihei et al., 2000) and long-term depression (LTD) (Zhao et al, 1999) in the HC, reduced HC spine density (Zou et al., 2023), and reduced entorhinal cortex inputs to the HC (Zou et a., 2023). This ultimately leads to HC-dependent cognitive impairments (Kuhlmann et al., 1997; Jett et al., 1997; Munoz et al., 1988; Jaako- Movits et al., 2005; McGlothan et al., 2008; Wang et al., 2016). It is unknown how Pb^2+^ impacts HC network dynamics to mediate the effects of cellular pathologies on cognitive functions.

Beyond the direct effects of Pb^2+^ neurotoxicity, early-life Pb^2+^ exposure is further implicated in the development of SZ (Opler et al., 2004; 2008). A current leading hypothesis proposes that Pb^2+^ exposure escalates the risk for SZ through interactions of neurotoxic mechanisms with genetic and epigenetic factors at critical developmental stages (Guilarte et al., 2012). Supporting this view, early-life Pb^2+^ levels of patients *diagnosed* with schizophrenia have been found to significantly exceed those of control subjects, and postmortem assays of confirmed SZ patients have shown selective elevation of Pb^2+^ levels (Modabbernia et al., 2016). Additionally, Pb^2+^ exposure worsens behavioral, cognitive, and sensorimotor dysfunctions in animal models of SZ (Abazyan et al., 2014).

Interestingly, at a cellular level, developmental Pb^2+^ exposure recapitulates a hallmark of schizophrenia pathology by causing a reduction in the number of parvalbumin-expressing GABAergic interneurons (PVGIs) in the HC and agranular mPFC (Hashimoto et al., 2003; Konradi et al., 2011; Stansfield et al., 2015). These fast-spiking PVGIs are a critical component of mechanisms underlying gamma oscillations (25-120 Hz) that coordinate neuronal assemblies. Gamma oscillations stem from reciprocal inhibitory-excitatory (I-E) interactions of principal neurons nested within the local microcircuitry of inhibitory networks. I-E interactions also give rise to the HC theta rhythm (6-10 Hz) that coordinates gamma oscillations nested within each theta cycle (Tamura et al., 2017). Misbalance in these network dynamics possibly accounts for comorbid evidence from electroencephalographic (EEG) showing systems-level Pb^2+^ neurotoxicity disruptions (Kmiecik-Malecka et al., 2009; Govoni et al., 1988). In young children, low levels of Pb^2+^ correlate to slow-wave activity in sensory-evoked EEG data and overall theta power (Otto et al., 1981; Poblano et al., 2001). Additionally, Pb^2+^ exposure may also relate to some forms of epilepsy (Sasmaz et al., 2003) and increased seizures (Arrieta et al., 2005), further implicating network-level physiological disruptions as a critical level of analysis for chronic early life Pb^2+^ exposure.

Here, we explored the effects of Pb^2+^ exposure on network activity in the HC. To this aim, we performed chronic *in vivo* recordings (silicon arrays) in a rodent model of chronic early-life Pb^2+^ exposure (CELLE)(see Albores-Garcia et al. 2021). We evaluated delta, theta, and gamma frequency synchronization in HC networks (Schultheiss et al. 2020). We show that Pb^2+^ exposure caused *hypersynchrony* of the HC theta rhythm, an amplification of gamma oscillations, a disruption in behavior-related theta and gamma, and an increase in SWD indicative of absence seizures. Next, we tested the same Pb^2+^ exposed rat model (males and females) in PPI, a sensitive sensorimotor gating measure dependent on HC functions and related to SZ. We found that only male rats had significantly impaired sensorimotor gating, consistent with the Pb^2+^ exposure-induced HC network disruptions, SZ, and recapitulating sex differences in schizophrenics tested on a human version of the PPI task (e.g., Mena et al., 2016). We propose that theta hypersynchrony, theta-gamma activity patterns, and absence seizures represent key network dysfunction caused by early-life Pb^2+^ neurotoxicity, and likely contribute to associated developmental neuropathologies, including schizophrenia and childhood-onset epilepsies.

## Methods

### Subjects

Eleven adult Long-Evans rats (males) were implanted with commercially available silicon probes (NeuroNexus) and used in the electrophysiological experiments. In one rat, the probe was located in the lateral ventricle adjacent to the HC and was excluded. In another rat, tetrode signals were poor, and the rat was also excluded, leaving nine rats with good implants. Five rats were chronically exposed to Pb^2+^ (see below), and four were controls. Four additional adult male Long-Evans rats were implanted with custom-built, dual-site implants for simultaneous recordings from the HC and mPFC. Two dual-site rats were chronically exposed to Pb^2+^, and two were controls. These rats were only used to look at the relationship, if any, between HC SWDs and PFC activity. Two hundred thirty Long-Evans rats were used in pre-pulse inhibition (PPI) experiments. These rats were evenly divided into twelve groups (18-21 per group) corresponding to all combinations of (1) Pb^2+^ exposure and controls, (2) males and females, and (3) three ages at the time of testing.

All rats were individually housed in a climate-controlled vivarium on a reversed 12hr light/dark cycle, and experiments were conducted during dark cycle active periods. Rats had free access to food and water in their home cages, but no food or water was available during behavioral sessions. Rats with dual-site implants, but not rats in the other groups, received rewards (∼¼ Fruit Loop) during behavioral sessions delivered sporadically to augment rats’ motivation for exploratory behavior.

Experiments were conducted per the Guide for the Care and Use of Laboratory Animals (National Institutes of Health) following protocols approved by Florida International University’s Institutional Animal Care and Use Committee (FIU IACUC). Data from the six control rats used here had been previously analyzed to establish the relationships of HC spectral modes to behaviors (Schultheiss et al., 2020). The present study focused on the effects of Pb^2+^ exposure, and all analyses presented here are new.

### Breeding and chronic Pb^2+^ exposure

Adult female rats (225-250g) (Charles River Laboratories) were randomly assigned to a Prolab RMH-1000 diet containing 0 (control) or 1500ppm lead acetate (PbAc) (Dyets, Inc., Bethlehem, PA). After 10-14 days of acclimation to the diet, each dam was paired with a non-exposed adult male Long Evans rat (250-275g; Charles River Laboratories) to breed over 3 days. Litters were culled to 10 pups on postnatal day one (PN1). Dams were maintained on their respective diet, and at PN21 pups were weaned onto the same diet and maintained for the duration of the experiments. Blood Pb^2+^ levels of rats used in PPI experiments (SuppTab1) were measured in samples obtained from cardiac puncture under sodium pentobarbital anesthesia. Samples were prepared and measured following protocols provided using the LeadCare Plus system (Magellan Diagnostics, N. Billerica, MA).

### Electrodes

For most electrophysiological experiments reported here, silicon probes with 32 electrodes arranged as eight tetrodes were surgically implanted to record LFPs from the dorsal HC. The eight tetrodes were distributed across four probe shanks at tip-to-tetrode depths of 78 µm and 228 µm (NeuroNexus A4X2-tet-5mm). Shanks were separated by 200 µm, giving each probe a total length of 0.67 cm, which was oriented medial-lateral when implanted to sample along the proximal-distal axis near the CA1 subregion of HC. Electrode impedances were measured to be 1.23 ± 0.32 MΩ (@1000Hz).

Custom-made dual-site implants housed 32 stainless steel wire electrodes (0.42 ± 0.25 MΩ) coupled to an integrated electrode interface board within a 3-D printed implant body (Autodesk Inventor; 3D Systems ProJet 1200). The electrode layout for these implants consisted of two grids, such that 18 electrodes targeted HC (2 x 2.5 mm grid, four electrode rows, 0.5 mm spacing), and 14 electrodes were aligned to the rostrocaudal axis of mPFC (4 x 1 mm grid, two rows, 0.5 mm spacing). HC sites were used in the primary analyses, while mPFC sites were used in supplemental analyses.

### Surgical procedures

Rats used in electrophysiological experiments weighed 600-700 g at the time of surgery. General anesthesia was induced with isoflurane (5%) mixed with oxygen (0.8 L/min) that was continuously delivered throughout surgery at 1-4% as needed. Body temperature was monitored throughout surgery, and a Ringer’s solution (5% dextrose) was given periodically to maintain hydration (1 ml increments every 60-90 minutes). Once the hair over the scalp was cut, rats were placed in a stereotaxic device, and glycopyrrolate (0.5 mg/kg, s.c.) was administered to assist respiratory stability.

Ophthalmic ointment was then applied to the eyes. Four injections of marcaine were made to the scalp (∼0.1 ml, 7.5 mg/ml, s.c.), and a rostrocaudal incision was made, exposing the skull. The pitch of the skull was leveled between bregma and lambda, two stainless steel ground screws were placed in the left parietal bone, and five titanium support screws were anchored to the skull. For silicon probe implants targeting dorsal HC, rectangular craniotomies were drilled to accommodate the probe shanks centered on coordinates AP -3.2 mm, ML 2.7 mm. For dual-site implants, craniotomies were shaped to accommodate the respective electrode grids targeting HC (AP -2.9 mm, ML 2.0 mm) and mPFC (AP 3.0 mm, ML 1.0 mm).

After removal of the dura, implants were lowered on the stereotaxic arm until the tips of the probe shanks (or electrode wires) were just above the cortical surface. Then, the ground wire was attached to the ground screws, and implants were lowered further until the electrodes reached a depth of ∼2.8 mm below the cortical surface. A thin layer of sodium alginate was applied in the craniotomy to protect the exposed surface of the brain, and dental cement was used to secure the implant to the skull screws permanently.

Neosporin^®^ was applied to the skin surrounding the implant, and flunixin (2.5 mg/kg, s.c.) was given for analgesia. Before being returned to their home cages after surgery, rats were allowed to rest on a heating pad until ambulatory. Neosporin^®^ and flunixin were given again on the day following surgery, and rats were monitored closely for five days to maintain body weight. Daily Baytril injections (s.c.) were also given for five days to protect against postsurgical infection. A minimum of one week was reserved for recovery before beginning experiments.

### Behavioral setup for electrophysiological recordings

All electrophysiological experiments were conducted in a large open-field environment (122 x 118 x 47 cm). A custom-made white cutting board served as the floor of the behavioral arena, and the interior walls were lined with white vinyl. The arena was mounted on posts 72 cm above the floor, and the exterior was wrapped in grounded Faraday shielding.

Two video cameras with a bird’s eye view were used for behavioral tracking. These were mounted 28 cm apart and 150 cm above the arena floor, providing ∼4.6 pixel/cm resolution in recorded videos and corresponding tracking data.

The entire behavioral apparatus was enshrouded with black curtains suspended from the ceiling and extending to the floor. Small gaps at the Velcro^®^ closures between the four curtain panels were used to observe rats covertly to monitor that the head stage, cables, and elastic tethers were not tangled and to confirm that rats were awake during periods of relative inactivity. All room lights were kept off during recording sessions, and the white arena was illuminated dimly with red light from LED strips mounted at the level of the video cameras above. During behavioral sessions, rats were placed on the arena floor and recorded without any intervention except for infrequent behavioral checks or to adjust cables or connections.

### Electrophysiological recordings

Throughout each experiment, wideband neural data were acquired with a Plexon OmniplexD recording system with A/D conversion at 40 kHz (16-bit). A digital head stage processing subsystem (Intan, 32 channel) passed continuous wideband data referenced against the ground screws (low cutoff = 0.7 Hz) to a hard drive using PlexControl data acquisition software. For analysis of LFP recordings, wideband data were downsampled to 2 kHz. A maximum of one channel per tetrode was included in our analyses, and several tetrodes across rats were excluded due to poor signal. With these restrictions, 30 channels recorded from Pb^2+^-exposed rats and 30 channels from control rats were included in our analysis of HC LFP power spectra during running and stationary behaviors. These 60 channels were selected from among the 288 total recorded channels using a custom algorithm designed to maximize signal quality by choosing for each tetrode the channel with the highest theta-delta ratio during running (t-d_run_) when theta would be expected to be strongest and delta weakest. Channel selection for analyses of SWD (detailed below) was done manually through the inspection of several basic features, such as the lack of evident noise in the voltage plots, the presence of separable theta and delta modes throughout the session, and the ability to separate SWD-related theta and theta harmonics and running- related theta and theta harmonics.

### Behavioral tracking

The physical and software settings for the two video cameras used for behavioral tracking were tuned independently to optimize detection of body contours on one camera and to optimize tracking of head-mounted LEDs on the other. Using Cineplex software (Plexon), tracking data and video frame captures (30 fps) were synchronized in real time to the simultaneously recorded neural data via a central timing board. LED tracking data were collected online during experiments, and body contour tracking data were derived offline from recorded videos.

LED tracking entailed tabulating the coordinates of two clusters of LED florets (green and blue) that were offset ∼2 cm laterally from the head stage at ∼3 cm above the rat’s head. Each LED cluster comprised three florets oriented upward and slightly outward. This construction enabled consistent LED tracking despite behaviors in which rats oriented their heads away from the level plane. The location of the head was calculated as the midpoint between the two LED clusters for each time point. Body location was calculated from difference images obtained by subtracting an image of the empty arena from each frame capture during an experiment. Movement speeds were calculated from these body coordinates, except that head location was used to calculate movement speeds in our seizure detection protocol (below).

### Co-registration of LED and body tracking data

Following behavioral data acquisition, all analyses were performed with custom-written MATLAB scripts (MathWorks, Natick, MA). First, the coordinate system for camera one was mapped onto the coordinate system for camera two, enabling the integration of the two types of tracking data. Next, for each timestep, we derived rats’ head direction (HD) relative to the environment (from the axis between the two head-mounted LED clusters); body direction (BD) relative to the environment (from the midpoint between the LEDs to the center of the body); head angle (HA) relative to the body (using the LED axis relative to BD); and the rotational velocity of the body (from changes in BD).

### Definitions of running and stationary bouts

Rats exhibit a wide range of behaviors in the open field, including frequent rearing and grooming behaviors that incorporate fast body and head movements. These movements manifest in tracking data as changes in body and LED coordinates, just as running does. To isolate bouts of locomotion from other behaviors, we first required that movement speed be maintained above 5 cm/sec for a minimum duration of 1.025 or 2.05 seconds (in parallel analyses) with a minimum peak speed of 15 cm/sec. These minimum bout durations were chosen as a compromise between the relatively fast timescale of rats’ locomotor behaviors (e.g., rapid transitions between brief periods of continuous running and interspersed pauses), the frequency resolution of power spectra, and the reduction in number of oscillatory cycles (particularly for delta) contributing to amplitude and power estimates from shorter time windows. We then eliminated all instances in which the head angle exceeded 35 degrees or body rotational velocity exceeded 52.5 degrees per second at any time. Stationary bouts were required to meet the same criteria for minimum duration, HA, and rotational velocity, and movement speeds were required to remain below 5 cm/sec.

### Measures of oscillatory synchronization

We analyzed delta-, theta-, and slow and fast gamma-frequency synchronization (1-4 Hz, 6-10 Hz, 25-55 Hz, and 65-120 Hz, respectively) using short-time power spectra (PSDs) of the HC LFP, as well as peak-to-trough measurements of individual oscillatory cycles of the respective band-pass filtered LFPs. We also derived a continuous time series of instantaneous amplitudes for each band by resampling cycle-by-cycle amplitude measures to 2 kHz. This transform was applied in our analyses of speed modulation and cross-frequency correlations to mitigate the vanishing likelihood that longer oscillatory cycles will elapse precisely within the time course of brief behavioral bouts.

### Boutwise analyses

We calculated the spectral content of HC recordings using Welch’s method for each stationary and running bout. Estimates of delta and theta power from PSDs for individual behavioral bouts were defined as the mean values across the respective frequency ranges. Delta and theta peak frequency and power estimates were measured directly from PSDs for behavioral bouts. For each recording site, delta and theta power were defined as the mean values across all bouts of each type. To determine whether Pb^2+^ exposure qualitatively impacts the behavioral modulation of power spectra, for each rat, we subtracted the mean PSD for stationary bouts from the mean running spectrum. These *difference* PSDs were then normalized either by the area of the mean stationary PSD, or by calculating the percent change at each frequency. We also compiled the combined distributions of cycle amplitudes for each frequency band across all rats in each group during all bouts of each type across all rats in each group.

### Sliding time-window analyses

To elaborate on bout-wise comparisons, we averaged movement speeds and continuous amplitude time series for each frequency band using a two-second sliding window (50% overlap) across each session for each rat. We then averaged oscillatory amplitudes within 5 cm/sec movement speed bins up to 75 cm/sec. This approach, which is typical for evaluating speed modulation of theta, ignores the distinction between running and movements during other behaviors, but it captures the full nonlinear relationships of synchronization in each frequency band across the continuous range of movement speeds. We calculated pairwise correlation coefficients between speed and amplitude measures for each frequency band, and between bands.

### Peak speed-triggered analyses

In a last series of analyses, we focused on the secondary regime of running speed modulation by triggering cycle-by-cycle amplitudes on local peaks in running speed (>40 cm/sec) separated by at least 10 seconds. We then averaged amplitude measures within 0.5-second bins spanning -9.75 to 9.75-second latencies. Because behavioral variability increases at greater temporal offsets from speed peaks, the fidelity of relationships between time-binned cycle amplitudes and running speed is maximal at the shortest latencies. We focused our initial statistical analyses on the 5 speed bins spanning the 2.5-second window from -2.25 to 0.25-second latencies. We compared unnormalized cycle amplitudes for each frequency band between Pb^2+^-exposed rats and controls. We z-normalized cycle amplitudes in two ways, in separate analyses, to address different effects of Pb^2+^ on synchronization. To balance the contributions of each recording site to group-wise comparisons, we z-normalized oscillatory amplitudes in each frequency band across the duration of each behavioral session. Next, we z-normalized amplitudes across each peak-triggered analysis window separately before averaging within bins. By eliminating the implicit assumption that amplitude measures are stable throughout behavioral sessions, this approach had the effect of drawing running speed modulation of amplitudes into greater relief. That is, changes in amplitude accompanying changes in speed were normalized relative to temporally local amplitudes only, revealing a novel and reliable pattern of slow fluctuations in theta and gamma synchronization that unfolded over the several seconds preceding and following speed peaks.

### Absence seizure detection

Although seizures are rare and highly variable across rats, sessions, and strains, rodents are a universal model for synchronized SWDs that dominate the LFP across the cortex during non-convulsive seizures in the absence of epilepsy (e.g., van Luijelaar et al., 2014). SWDs occurred overwhelmingly at times when rats were stationary, supporting the view that SWDs are the physiological manifestations of ’absence’ seizures. We estimated the incidence and duration of seizures by the experimenter’s hand measurements. Seizure parameters differed between the Pb^2+^-exposed and control groups, but biased judgment could manufacture the same effects. We developed a novel seizure detection algorithm based on the frequency composition of the short time-windowed LFP to (1) obtain unbiased estimates of the incidence of SWDs, and to (2) compare seizure parameters between Pb^2+^-exposed rats and controls. Using windows of two seconds (one-second overlap), we first derived power estimates within a 2 Hz frequency range centered on the frequency of peak theta power (5-14 Hz) and within 2 Hz ranges centered on the first and second harmonics of the theta-peak frequency. We took the z-scored values across all time windows for each frequency band throughout each session. Negative values in these time series were zeroed, yielding metrics of elevated theta and theta harmonic power. These metrics and the time-averaged LFP amplitude were smoothed with a Gaussian, and their product was z-scored to give a timeseries that was effective at identifying most SWDs of reasonable strength. We categorized detection peaks as SWDs when they exceeded threshold values of 2 and only when movement speeds were below 5 cm/sec within the corresponding PSD windows. The incidence, strength, and duration of SWDs were then compared between Pb^2+^-exposed rats and controls.

### PPI task

Pre-pulse inhibition of acoustic startle (PPI) was assessed in male and female rats at PN28, PN50, and PN120 following a protocol adapted from Abazyan et al. (2014). Four identical SR-LAB startle chambers (San Diego Instruments Inc., San Diego, CA, USA) were used to measure startle responses. Rats were placed inside the startle chamber into a mounted Plexiglas® cylinder with an accelerometer. Broadband background noise and acoustic stimuli were delivered via a speaker on the chamber’s ceiling. Sound levels were calibrated using a digital sound level meter, and accelerometer sensitivities for each chamber were calibrated daily. The SR-LAB software and interface system controlled the presentation of acoustic stimuli, and accelerometer measurements were recorded. Each experimental session started with a five-minute period during which rats acclimatized to 68 dB of background noise that was continuous throughout the session. Rats’ average startle responses were calculated as the mean of 10 acoustic pulses (120dB, 100ms) presented at random intertrial intervals. Rats were left in the enclosure for an additional five minutes before beginning PPI trials.

Each rat was exposed to the following trial types: pulse-alone (120 dB, 100 ms); pre-pulse alone (90 dB, 20 ms); and pre-pulse–pulse combinations using one of five pre-pulse intensities: 74, 78, 82, 86, and 90 dB. Pulse onsets lagged pre-pulse offsets by 80 ms. Each session consisted of six trials with each pre-pulse intensity, presented in a pseudorandom order with random inter-trial intervals. Startle responses were quantified as accelerometer measurements averaged over the 100 ms pulse presentations. PPI was quantified as the percent change in startle amplitude on pre-pulse–pulse trials relative to pulse-alone trials.

### Histology

After data collection, rats used in electrophysiological experiments were anesthetized with isoflurane (5%) mixed with oxygen (800 ml/min), and marking lesions were made with a NanoZ (Plexon) to deliver 10μA for 10s at each electrode location. An hour later, rats were transcardially perfused with 100 ml phosphate-buffered saline (PBS), followed by 200 ml of 4% paraformaldehyde (PFA, pH 7.4; Sigma-Aldrich, St. Louis, MO). Blood Pb^2+^ levels of these rats were measured in samples obtained from cardiac puncture during the perfusion. Brains were post-fixed overnight in 4% PFA and then placed in a 30% sucrose solution for cryoprotection. Frozen brains were cut on a Leica CM3050 S cryostat (40 μm; coronal) and Nissl-stained. Images were acquired on an Olympus BX41 with a UPlanSApo 4x/0.16 objective under brightfield illumination. Modest brightness, color, and contrast adjustments were made uniformly in macOS Sonoma 14.5 using Preview. Lesion locations were determined by multiple raters using the microscope directly, and then matched to the photomicrographs, which were marked using plates from the Paxinos & Watson Atlas (2004).

## Results

### Electrophysiological recording sites

In nine rats, NeuroNexus probes were localized in the dorsal HC, primarily in the stratum radiatum (rad) and stratum lacunosum (slm) layers of CA1, with some electrode markings spanning the CA1-CA2-CA3 transition zone, and one electrode was deeper near DG (SuppFig1A). In four rats, the implanted stainless-steel wire probes had electrode markings in the prelimbic region (PL) of mPFC and the HC, mostly the slm layer of CA1, with some wires encroaching on CA2/CA3 subregions (SuppFig1B). For these probes, only medial sites in the mPFC were utilized. SupplFig1 shows the histology for all rats.

### Blood Pb^2+^ levels (BLLs)

Post-mortem BLLs were more than 12-fold higher in the Pb^2+^ exposed electrophysiology rats (five rats sampled: 29.7, 30.3, 38.3, 38.4, and 41.3 μg/dL) compared to controls (two rats sampled: 2.8 and 2.8 μg/dL). These BLLs are at the higher end of the range typically observed in the 1500ppm CELLE paradigm, likely because these rats were much older than normally used at ∼11 months. SuppTab1 shows the BLLs for the PPI rats in all groups (ages PN28, PN50, and PN120). In the PPI Pb^2+^ rats, BLLs were between 19 and 25 μg/dL, and the controls were all less than 1.9 μg/dL (limit of detection).

### Grossly intact modes of HC synchronization and exploratory behavior in Pb^2+^ exposed rats

To characterize any HC pathophysiology caused by chronic Pb^2+^ exposure, we evaluated the effects of running and stationary modes on oscillatory synchronization in the HC. We first assessed the gross spectral content of HC LFPs as rats freely explored a large ’open field’ environment Fig1 A-D).

Spectrograms illustrating the time-frequency decomposition of HC LFPs were visually similar between exposure groups (Fig1E and F). In both Pb^2+^-exposed rats and controls, there were stronger theta rhythms during running (green arrows) and elevated delta-band power when rats were stationary (cyan arrows).

This observation suggests that chronic Pb^2+^ exposure does not disrupt HC networks so severely as to subvert the fundamental modes of oscillatory synchronization that coordinate neuronal ensembles and cortico-HC circuit dynamics during behavioral episodes (Hyman et al., 2005; Jones & Wilson, 2005; Siapas et al., 2005; Sirota et al., 2008; van der Meer & Redish, 2011).

Importantly, we did not observe any effects of Pb^2+^ exposure on several continuously sampled measures of rats’ locomotor behaviors in plots of continuous movement speed, spatial orientation (body and head direction), and rotational velocity (SuppFig2); nor did we find differences in behavioral measures for distinct bouts of running (number of bouts: t_(7)_ = 0.271, p = 0.979; speed: t_(7)_ = 0.186, p = 0.858; and duration: t_(7)_ = 0.172, p = 0.868), or during stationary bouts (number of bouts: t_(7)_ = 0.261, p = 0.801; speed: t_(7)_ = 0.912, p = 0.392; duration: t_(7)_ = 1.806, p = 0.114)(SuppFig2). These observations suggest no gross changes in open-field behavior following Pb^2+^ exposure, which might have directly affected electrophysiological activity.

### Absence seizures in Pb^2+^-exposed rats and controls

In addition to quasi-normal delta- and theta-dominated network modes coupled to the locomotor stops and starts, during quiet immobility, we observed sporadic instances of maximal theta synchronization accompanied by striking theta harmonics in both Pb^2+^ (Fig 1F) and control rats (Fig 1E). LFP waveforms corresponding to these theta-harmonic events were typical of SWDs and reflective of the neurophysiological manifestations of ’absence seizures’ (Fig 1F)(Jafarian et al., 2020). SWD incidence was highly variable across rats and between recording sessions for individual rats, but all rats exhibited SWD during at least one session, in line with previous reports (e.g., Taylor et al., 2019; Rodgers et al., 2015; Letts et al., 2014; Pearce et al., 2014).

**Figure 1.**
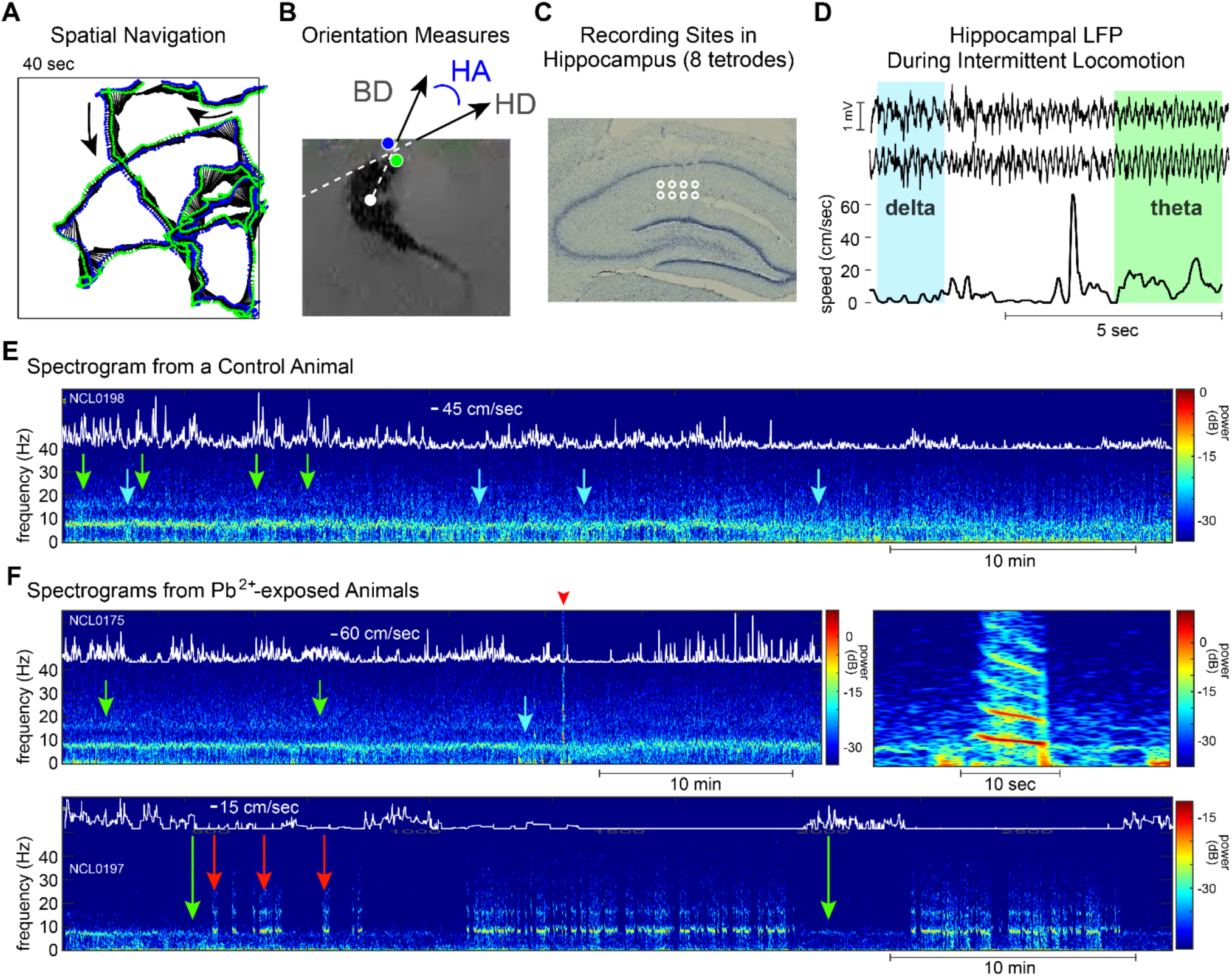
During exploratory navigation, HC LFPs display grossly similar spectral content in Pb^2+^-exposed rats as controls. **A.** Representative spatial trajectory during free exploration of the square (120 cm) ’open field’ environment. Black lines represent the rat’s body direction (BD as in **B**), and green and blue dots represent head-mounted LEDs. **B**. Continuous measures of locomotor behavior, including head direction relative to the environment (HD) and head angle (HA, derived from HD and BD). **C**. Recording sites of 8 tetrodes located at two depths on each of four shanks of a silicon probe. Tracks left by the shanks of the implanted probe are visible as disruptions of the CA1 cell layer. **D.** Representative LFP traces (top) during running and immobility (bottom) showed corresponding periods of strong theta and delta-band activity (e.g., as highlighted in cyan and green). **E & F**. Representative spectrograms for control rats (**E**) and Pb^2+^-exposed rats (**F, top left panel**) showed qualitatively similar patterns of speed modulation (overlaid white traces) of theta (green arrows) and delta (cyan arrows). **F, top right panel**. Putative absence seizures occurred in all Pb^2+^-exposed rats. Seizure incidence was highly variable across rats and sessions for each rat, i.e., the spectrograms shown in **F** illustrate sessions during which a single seizure lasting 7 seconds and ∼25 minutes of continuous seizing occurred (**top** and **bottom panels, respectively)**. The red arrowhead in the **top left** panel indicates the seizure magnified in the **top right**. Green arrows (**bottom panel**) indicate running accompanied by elevated theta power. Red arrows indicate seizure bouts during pauses in the rat’s intermittent locomotor behaviors.

### Theta rhythm hypersynchrony

Power spectra of HC LFPs confirmed that awake delta and theta rhythmic network modes were superficially intact in Pb^2+^-exposed rats (Fig 2A), although there was substantial variability across tetrodes. Representative PSDs across all eight tetrodes for a lead and control rat are illustrated in Fig 2A. Next, we ran a more detailed analysis, restricting PSD measures to periods of running and stationary behaviors. Using this approach, we found that theta synchronization was strikingly elevated in Pb^2+^-exposed rats (30 channels) compared to controls (30 channels) during bouts of running (2- sec windows: t_(58)_ = 2.786, p = 0.007; 1-sec windows: t_(58)_ = 3.177, p = 0.002) and stationary behaviors (2- sec windows: t_(58)_ = 2.056, p = 0.044; 1-sec windows: t(58) = 2.632, p=0.011) (Fig2B). Moreover, theta power (dB: t_(58)_ = 2.756, p = 0.008; %: t_(58)_ = 1.711, p = 0.093, a significant trend) and theta-harmonics (dB: t_(58)_ = 3.125, p = 0.003; %: t_(58)_ = 3.106, p = 0.003) were also greater in Pb^2+^-exposed rats compared to controls during running, measured relative to stationary behaviors (Fig2C). Altogether, this suggests stronger theta rhythmogenic mechanisms and/or stronger activation of those mechanisms in HC networks in Pb^2+^ exposed rats. This also suggests that Pb^2+^ exposure-based augmentation of the difference in theta synchronization between behavioral states may reflect an increased propensity of HC network activity to self-organize theta rhythmicity given appropriate behavioral excitation, or it may reflect increased susceptibility of HC to entrainment by theta rhythmic inputs recruited during running.

**Figure 2.**
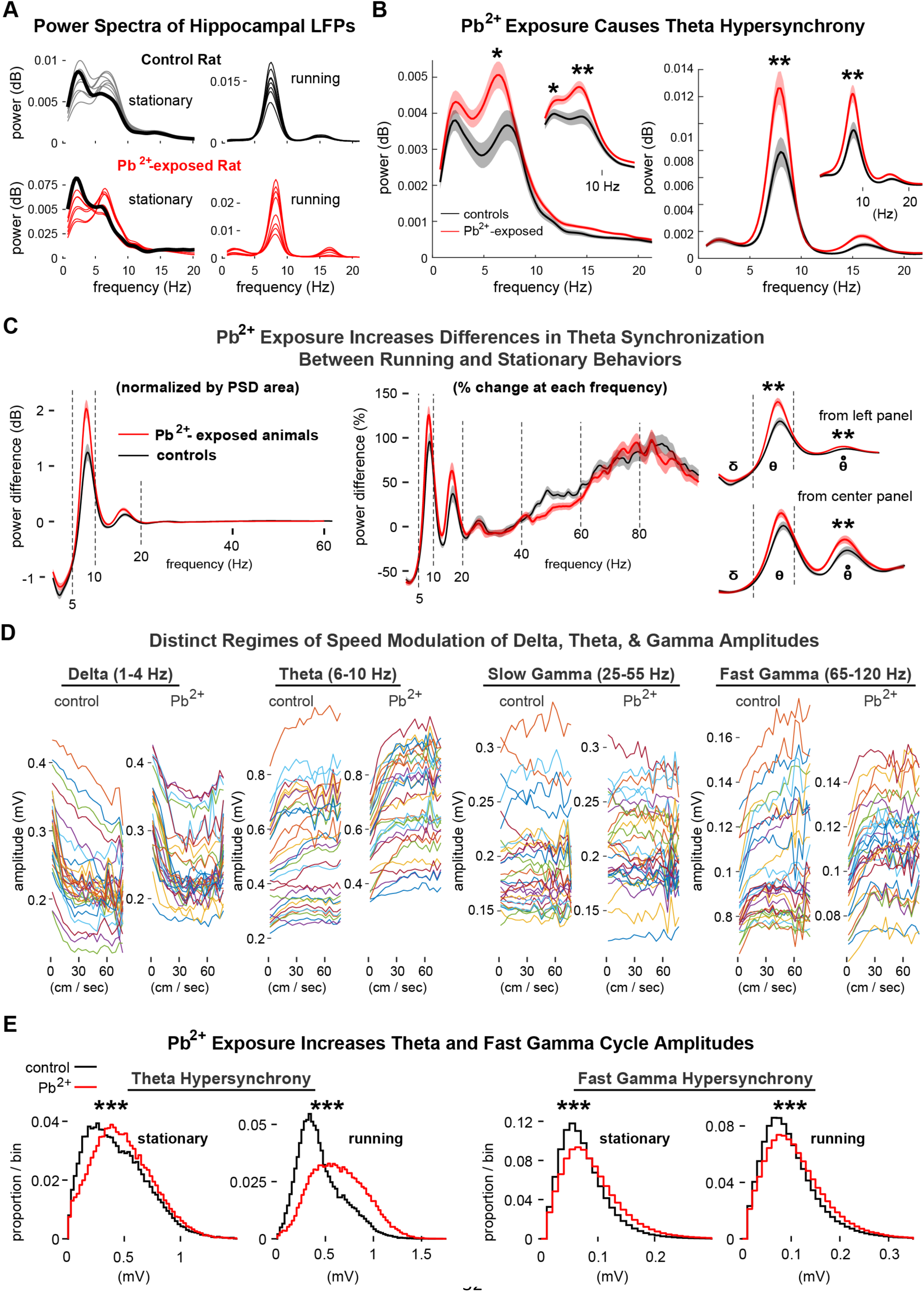
Theta-frequency hypersynchrony occurs in rats exposed to Pb^2+^. **A.** Power spectral densities (PSDs) of HC LFPs both for Pb^2+^-exposed rats and for controls exhibited diminished delta and heightened theta during running (**right**) relative to stationary bouts (**left**). Grey and red traces indicate representative PSDs of simultaneous recordings from each site in the HC in individual rats. Thick black traces (**left panels**) are PSDs from the site with the highest delta power during immobility, highlighting the apparent tradeoff between delta and theta peaks across recording sites in each animal. **B**. Marked elevation of theta power in Pb^2+^-exposed rats relative to controls during two-second bouts of immobility (**left**) and running (**right**) (1 sec bout **insets**). **C**. Stronger behavioral modulation of theta power in Pb^2+^- exposed rats than controls in terms of the absolute change (normalized by total power (0-25 Hz) (**left**) and the relative change at each frequency (%) (**right**) between stationary and running bouts for control (black) and Pb^2+^-exposed rats (red). Enlargement of the low-frequency range of PSDs in the left and center panels (**right**). **D.** Running speed modulation of delta, theta, slow, and high gamma. **E**. Similar patterns of movement speed modulation of cycle-by-cycle amplitudes of theta and gamma oscillations (across all behaviors) in Pb^2+^-exposed rats and controls. Significance is indicated as follows: *, p<0.05; ** p<0.01; ***, p<0.001.

Delta power in Pb^2+^-exposed rats was also elevated during 1-sec stationary bouts (t_(58)_ = 2.047, p = 0.045) (Fig 2B, inset), but this was neither significant nor a trend for 2-sec bouts (t_(58)_ = 1.488, p = 0.142) (Fig 2B). Delta power also showed a significant trend of elevated levels in Pb^2+^ rats relative to controls during running bouts of 1-sec windows, but not 2-sec windows (1-sec: t_(58)_ = 1.798, p = 0.077; 2-sec: t_(58)_ = 0.240, p = 0.812). There were no significant differences in delta between Pb^2+^ and control rats during running measured relative to stationary behaviors (dB: t_(58)_ = 0.374, p = 0.710; %: t_(58)_ = 0.7850, p = 0.436)(Fig 2C). Consistent with previous reports (Sakata et al., 2005), we also observed elevated delta power at the onset or offset of SWDs (SupplFig3). This activity could reflect some aspect of behavior that may be related to the onset of or escape from behavioral and cognitive arrest accompanying absence seizures, or it may reflect a behavior-independent neurophysiological signature that is coupled to transitions between seizure and non-seizure network states. Interestingly, linear fits to delta power across individual recording sites from each rat further revealed a consistent distal-proximal gradient of increasing delta power in control rats that was not evident in Pb^2+^-exposed rats (SuppFig4).

### Oscillatory amplitudes

Whereas delta or theta synchronization typically dominates the power spectrum, gamma frequency synchronization is not well captured by power estimates taken from PSDs.

Gamma power is diminutive relative to slower frequencies, in part because gamma synchronization can occur across a broad range of frequencies, while faster dynamics of power and frequency are generally averaged across longer windows, and the network dynamics underlying gamma oscillations can vary in scale from markedly localized network motifs to large-scale coherent circuits. Therefore, to evaluate the impact of Pb^2+^ on gamma synchronization and to further explore delta and theta effects, we derived peak-to-trough amplitudes of individual oscillatory cycles for each frequency band throughout each recording session.

### Modulation of oscillatory amplitudes by rats’ movement speeds

Cycle amplitudes for delta, theta, and fast gamma were related to movement speeds in control rats and Pb^2+^-exposed rats (Fig 2D). Each of these relationships comprised: (1) a steep primary regime at slow movement speeds (∼0-15 cm/sec) mainly reflecting averaging across transitions between network modes at behavioral transitions between immobility and running, and (2) a flatter secondary regime reflecting genuine running speed modulation of oscillatory amplitudes. To demonstrate the relationships between amplitudes and speeds, we ran correlations on each channel, counted the number of significant correlations across bands and groups, and the mean Pearson’s r as a measure of correlation strength. Results show that delta was negatively related to running speed in 21/30 Pb^2+^-exposed rats (mean r = -.508) and 29/30 controls (mean r = −0.727), primarily reflecting a quick drop in the amplitude during the initial regime. Theta was positively correlated with running speed in all 30/30 Pb^2+^-exposed rats (mean r = 0.544) and 23/30 controls (mean r = 0.739).

Comparatively, slow gamma had very weak and inconsistent (positive and negative) relationships with running speed, showing significant correlations in 9/30 Pb^2+^-exposed rats (mean r = -.090) and 14/30 controls (mean r = −0.093). Lastly, fast gamma amplitudes had strong positive relationships with running speeds, showing 21/30 Pb-exposed rats (mean r = 0.532) and 20/30 controls (mean r = 0.502).

### Pb^2+^ amplifies delta, theta, and fast gamma synchronization

Theta cycle amplitudes, compiled across rats in each group, were strikingly elevated in Pb^2+^-exposed rats relative to controls during stationary bouts (t_(114,635)_ = 32.920, p = 1.45x10^-236^) and running bouts (t_(102,653)_ = 116.521, p < 0.001) (Fig 2E). Fast gamma-frequency oscillations were also markedly amplified during stationary bouts (t_(1,276,491)_ = 168.199, p <0.001) and running (t_(1,127,377)_ = 101.326, p < 0.001)(Fig 2E). Interestingly, theta frequency was substantially reduced in Pb^2+^-exposed rats during stationary bouts (control: 7.159 ± 1.099 Hz, Pb: 6.837 ± 0.984 Hz; t_(842)_ = 4.489, p = 8.153 x 10^-6^) and running (control: 8.188 ± 0.439 Hz, Pb: 7.759 ± 0.602 Hz; t_(702)_ = 10.769, p = 3.9 x 10^-25^), potentially reflecting synchronization of larger neuronal populations. Differences in delta-frequency cycle amplitudes between Pb^2+^-exposed rats and controls were small relative to those for theta and gamma (stationary: t_(41,741)_ = 5.547, p = 2.932 x 10^-8^; running, t_(34,211)_ = 8.235, p = 1.860 x 10^-16^). It is also important to consider that with such large numbers of individual cycle amplitude measurements for each frequency band, statistical significance may not be all that informative. However, these results generally show that Pb^2+^ increases cycle amplitude in the delta, theta, and fast gamma bands, with the most substantial differences in the latter two bands (Fig 2E).

### Pb^2+^ exposure tightens oscillatory fidelity to behavior

Pairwise scatterplots of theta power, theta frequency, and running speed for each bout type suggested that (1) covariation of theta power and frequency was disrupted in Pb^2+^-exposed rats relative to controls, and (2) running speed modulation of theta frequency was confined to a narrower range of speeds; whereas (3) running speed modulation of theta power occupied similar parameter spaces in Pb^2+^-exposed rats and controls (SuppFig5).

Comparisons of distributions of correlation coefficients confirmed the expected positive correlations of theta power and frequency to running speed for most recording sites in both Pb^2+^-exposed rats and controls (SuppFig6). Importantly, the correlation of theta power to running speed was elevated in Pb^2+^-exposed rats relative to controls (t_(58)_ = 2.582; p = 0.012), potentially reflecting the weakening of HC processes underlying spatial or mnemonic representations in favor of entrainment or tuning to locomotor parameters.

Correlations between running speed and theta frequency were similar in strength between groups (t_(58)_ = 1.466; p = 0.178) despite the apparent restriction of speed modulation of theta to a narrower range of speeds in Pb^2-^-exposed rats (SuppFig5).

### Behavioral modulation of delta, theta, and gamma oscillations

We have shown that Pb^2+^-exposed rats exhibited theta rhythm hypersynchrony and higher amplitude gamma frequency oscillations relative to controls during stationary and running bouts, as well as increased correlations of oscillation parameters to locomotor behavior (Fig 2). To determine whether theta and gamma hypersynchrony during running were related to functional differences in speed modulation of oscillatory amplitudes, we evaluated the amplitudes of delta, theta, and fast gamma oscillations before and after peaks in running speed (>40 cm/sec) that were separated by at least 10 seconds. Average peak-triggered amplitudes of delta, theta, and fast gamma oscillations were elevated in Pb^2+^-exposed rats relative to controls throughout the peak-triggered analysis window (Fig 3A). The non-zero average values of these z-scored amplitudes stem from the elevated likelihood that an animal is running at times within the analysis window (recent or upcoming running occurring within +/- 10 sec) relative to more remote times. As predicted by our bout-wise analyses of oscillation amplitudes, delta cycle amplitudes were at their lowest near the times of running speed peaks.

**Figure 3.**
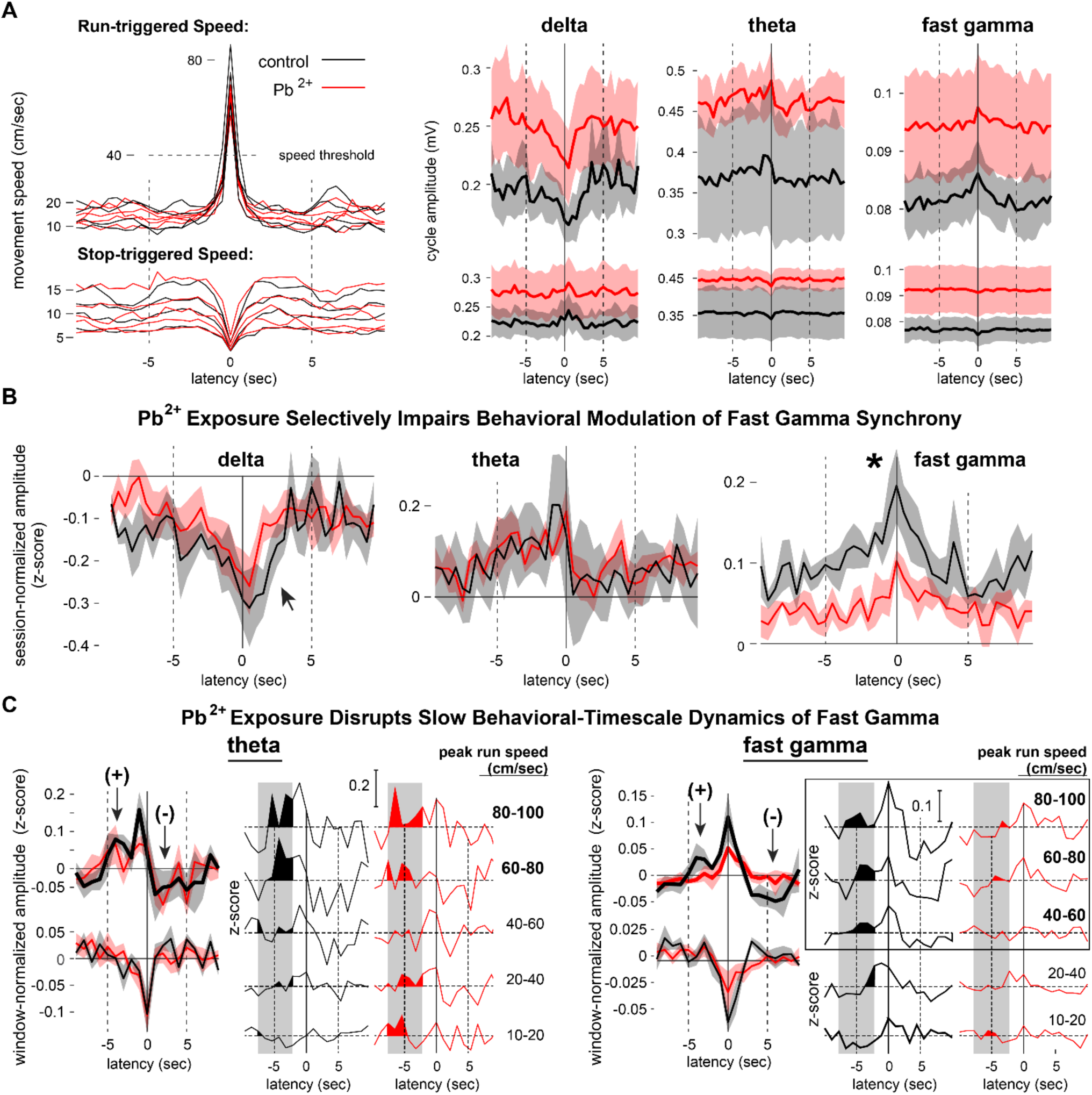
Pb^2+^ exposure leads to disruptions in behavior-triggered high gamma activity. **A.** Running- triggered speed- and stop-triggered speed transition for delta, theta, and high gamma amplitudes**. B.** Z- normalized amplitudes for each frequency band across the 20-second triggering window for each peak in running speed, emphasizing temporal dynamics disrupted at high gamma frequencies. **C.** Peak running speed separated in 20 cm/sec bins, showing fast gamma disruptions (right-most red line plots). Significance is indicated as follows: *, p<0.05.

In contrast, theta and fast gamma amplitudes were highest at or just preceding peaks in running speed (Fig 3A). The opposite pattern occurs with stop-triggered amplitudes (Fig3A).

### Pb^2+^ selectively disrupts the behavioral modulation of fast gamma

We asked whether Pb^2+^ exposure impacted the behavioral modulation of delta, theta, and fast gamma synchronization by comparing z-normalized amplitudes in each frequency band at short latencies from running speed peaks. Normalization across the duration of each recording session generated vectors representing the proportionate change in oscillation amplitude with the behavioral parameter. It also balanced each channel’s contribution to group-wise comparisons despite possible differences in the amplitude or quality of raw LFP signals. Fig 3A shows speed peak-triggered averages of relative amplitudes for delta, theta, and fast gamma. Normalized delta amplitudes were negative at all peak-latencies for both Pb^2+^-exposed and control rats, indicating that delta amplitudes near running speed peaks were smaller than the average across all behaviors (Fig 3B). In contrast, normalized amplitudes of theta and fast gamma oscillations were positive at all latencies, indicating elevated synchronization during fast running that, for theta, returned to near average levels immediately after speed peaks (Fig 3B), but for fast gamma, returned to average levels more slowly over several seconds. Using mixed-design ANOVAs, we compared normalized amplitudes of the delta, theta, and fast gamma oscillations during the five-time bins spanning -2.25 sec and 0.25-sec latencies between Pb^2+^-exposed and control rats. We found a main effect of decreasing delta amplitudes across latencies relative to speed peaks (F_(4, 44)_ = 3.752, p = 0.015), but there was no effect of Pb^2+^ exposure on delta synchronization relative to controls (F_(1, 44)_ = 0.194, p = 0.673), nor an interaction effect (F_(4, 28)_, = 0.174, p = 0.950). Interestingly, in this analysis, we found no difference across latencies for theta amplitudes (F_(4, 44)_ = 0.028, p = 0.872), no difference between Pb^2+^-exposed and control rats (F_(1,44)_ = 0.906, p = 0.474), and no interaction effect (F_(4, 28)_ = 1.851, p = 0.147). Notably, the ANOVA for fast gamma revealed significant main effects of Pb^2+^ exposure (F_(4, 44)_ = 7.119, p = 0.0321), latency (F_(4, 44)_ = 6.684, p = 6.589 x 10^-4^), but no interaction effect (F_(4, 28)_ = 0.485, p = 0.746) indicating that whereas unnormalized fast gamma oscillations were amplified in Pb^2+^-exposed rats (Fig 3A, right panel), the degree to which fast gamma amplitude was modulated by running speed was diminished relative to controls (Fig 3B, right panel). Both effects can be seen in Fig 3A, noting that although the raw amplitudes were larger in Pb^2+^- exposed rats, the range across which gamma amplitude varied as a function of peak latency was compressed relative to smaller, more broadly modulated amplitudes in control rats.

### Pb^2+^ disrupts slow dynamics of fast gamma

Lastly, we independently z-normalized amplitudes for each frequency band across the 20-second triggering window for each peak in running speed included in the analysis. This change to the normalization used in the previous analysis removes the assumption that the range and scale of oscillatory amplitudes are stationary for a given frequency band throughout a behavioral session. This analysis, more sensitive to temporal dynamics, revealed reliable amplifications of theta and fast gamma oscillations at ∼-4 sec latency from running speed peaks (Fig 3C) that could reflect behavioral or cognitive regularities occurring before a period of running. We asked whether chronic Pb^2+^ exposure disrupted these slow temporal dynamics of theta or fast gamma amplitudes during running and found that the elevation of gamma amplitude was selectively eliminated in Pb^2+^-exposed rats while the corresponding amplification of theta was preserved (Fig 3C). We considered whether amplifying theta and fast gamma oscillations before running speed peaks might be a spurious observation. To the contrary, however, we consider its presence in both frequency bands and argue for it being a real effect. We also argue that unless the effect were real, it would not be expected to show a relationship to running parameters. However, we found that the early peaks in theta and fast gamma amplitudes predominantly occurred in trials with fast peak speeds, supporting the possibility of cognitive activity or behavior that reliably precedes fast-running peaks.

### Pb^2+^ exposure exacerbates absence seizures

As noted earlier, both Pb^2+^-exposed rats and controls exhibited absence seizures while stationary. These events were characterized by SWDs in the LFP (Fig 4A), accompanied by maximal theta-frequency synchronization and several theta harmonics appearing in spectrograms (Fig 4B). In many sessions, seizures generated a considerable right tail in distributions of theta power estimates during immobility (Fig 4C). In pronounced cases theta power distributions were bimodal (Fig 4C, inset) such that: (1) the lowest theta values corresponded to quiet-wakeful immobility; (2) intermediate and larger values corresponded to elevated theta during running; and (3) theta values during SWDs were on the order of the maximal theta during fast running (Fig 4D, left panel). Recordings at sites exhibiting weaker theta during running often enabled easy detection of SWDs (Fig 4D, right panel) using a single threshold for theta power. Including additional terms for thresholding power at theta harmonic frequencies further enabled the detection of SWDs (Fig 4D, right, insets).

**Figure 4.**
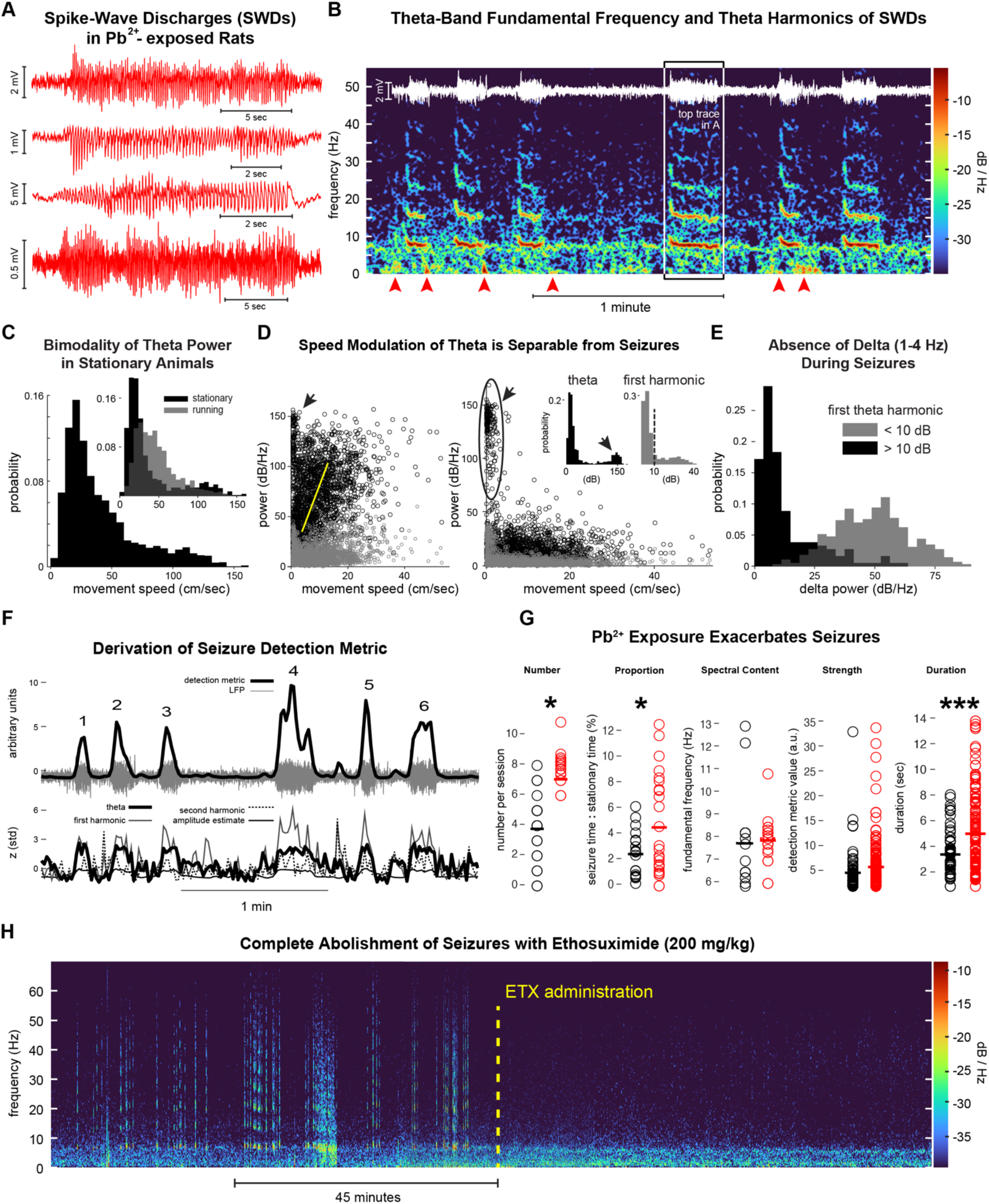
Chronic Pb^2+^ exposure exacerbates absence seizures in the HC. **A.** SWD (SWDs) in the HC LFP from four Pb^2+^-exposed rats (see SupplFig3 for coincident mPFC SWDs). **B**. Spectrogram showing six seizures. SWD overlaid in white. **C**. Theta power distributions during immobility show evidence of numerous seizure events creating a pronounced right tail or second peak at high theta power values. **D**. The theta power to movement speed relationship shows cases where theta during seizures corresponds to maximal theta during fast running (left). In other cases, weaker theta draws into relief the theta values corresponding to SWDs (right). Inset distributions of theta and harmonic power, as in C. **E**. Delta synchronizations, were minimal when power at the first harmonic of theta was elevated. **F**. Derivation of the seizure detection algorithm. **G**. The theta fundamental frequency did not differ between groups, but Pb^2+^- exposed rats exhibited exacerbated seizure number, proportion of stationary time, and duration. There was a trend toward significance in the strength of the seizures (p = 0.073). **H,** Sample spectrogram from one Pb^2+^ rat treated with ethosuximide (200mg/kg, i.p.) eliminates seizure but not normal theta and delta dominated running and stationary behaviors, respectively. Significance is indicated as follows: *, p<0.05; ***, p<0.001.

We recently characterized delta-dominated HC network activity during intermittent locomotion as a fundamental mode of encoding or segmenting events within a behavioral episode (Schultheiss et al., 2020). Seizures were uniformly distinct from these delta states during immobility, exhibiting minimal delta power (Fig 4E). Interestingly, in some of our data, strong delta synchronization was observed to flank seizures in the spectrogram, occurring at seizure onset or offset (Fig 4B, red arrowheads), which could aid seizure prediction in a clinical setting.

Given the variability in the spectral content of recordings at different HC sites (as in Fig 4D, left vs right), discerning the spectral signature of SWDs could be a greater or lesser challenge. Using a novel seizure-detection algorithm that integrates the self-normalized (z-scored over time) power at theta frequency and at the first two harmonics of theta with the concurrent LFP signal amplitude, we were able to detect high-reliability SWDs occurring during immobility (Fig 4F), see Methods for algorithmic details. We would argue these are likely to have been absence seizures. We found a broad range of values for which the detected events could be visually confirmed by varying the detection threshold applied to this algorithm. The range across which the algorithm performed well was bounded below by false positives, where the algorithm identified other instances of theta harmonic activity, typically during running, in addition to SWDs. We reduced most false positives by requiring the animal to be stationary during the theta-harmonic event. The algorithm’s functional range was bounded above by false negatives, where the algorithm exclusively identified SWDs but missed some proportion of the true incidence. Pb^2+^-exposed rats exhibited *normal* SWDs with the same theta fundamental frequency as controls (t_(42)_ = 0.027, p = 0.979). However, Pb^2+^ exposed rats had significantly more SWDs (t_(42)_ = 2.428. p = 0.020), trended toward greater strength (t(247) = 1.801, p = 0.073), had significantly longer durations (t_(247)_ = 4.294, p = 2.518 x 10^-05^), and occurred during a significantly greater proportion time spent stationary (t_(42)_ = 2.162, p = 0.036)(Fig 4G). To further confirm the validity of these SWDs as seizure-like events, we used ethosuximide (200 mg/kg) injections part of the way through a session in some of the rats. We observed that ethosuximide eliminated the appearance of SWDs at doses that did not alter gross behavior, supporting that these events are absence seizures (Fig 4H).

### A schizophrenia-like pattern of sensorimotor gating deficits caused by Pb^2+^ exposure

The electrophysiology experiments predict that disrupted HC network hypersynchrony should disrupt HC-related cognitive tasks. To examine this possibility, we measured PPI of the acoustic startle response (ASR), a test of sensorimotor gating that is disrupted by HC damage (Pouzet et al., 1999; Bast et al., 2001a; 2001b; 2003), and a canonical dysfunction linked to schizophrenia. Chronic Pb^2+^ exposure caused significant impairments of PPI at all stimulus levels for both late adolescence (PN50) and young adults (PN120) males (Mixed ANOVA: dB Levels: F_(3.12, 109.23)_ = 53.49, p < 0.001; Pb^2+^ vs. controls: F_(1,35)_ = 11.32, p = 0.002; Interactions: F_(3.12, 109.23)_ = 2.56, p = 0.069) and PN120 (Mixed ANOVA: dB Levels: F_(2.88, 115.11)_ = 59.59, p < 0.001; Pb^2+^ vs. controls: F_(1,40)_ = 9.30, p = 0.004, Interaction: F_(2.88, 115.11)_ = 2.00, p = 0.12)(Fig 5), but neither juveniles at PN28 (Mixed ANOVA: dB Levels: F_(2.57,89.83)_ = 68.69, p < 0.001; Pb^2+^ vs. controls: F_(1,35)_ = 0.54, p = 0.47; Interaction: F_(2.57,89.83)_ = 2.56, p = 0.069) nor females at any age exhibited any deficits (Pb vs. Control p’s > 0.10; Fig 5)(SuppFig7). This pattern recapitulates the sex dependence of PPI deficits exhibited in male and female SZ subjects (Mena et al., 2016; Takahashi et al., 2008; Ochoa et al., 2012), supporting the view that our rodent model for studying the Pb^2+^ induced HC pathophysiology may also be relevant for understanding aberrant HC network dynamics in SZ.

**Figure 5.**
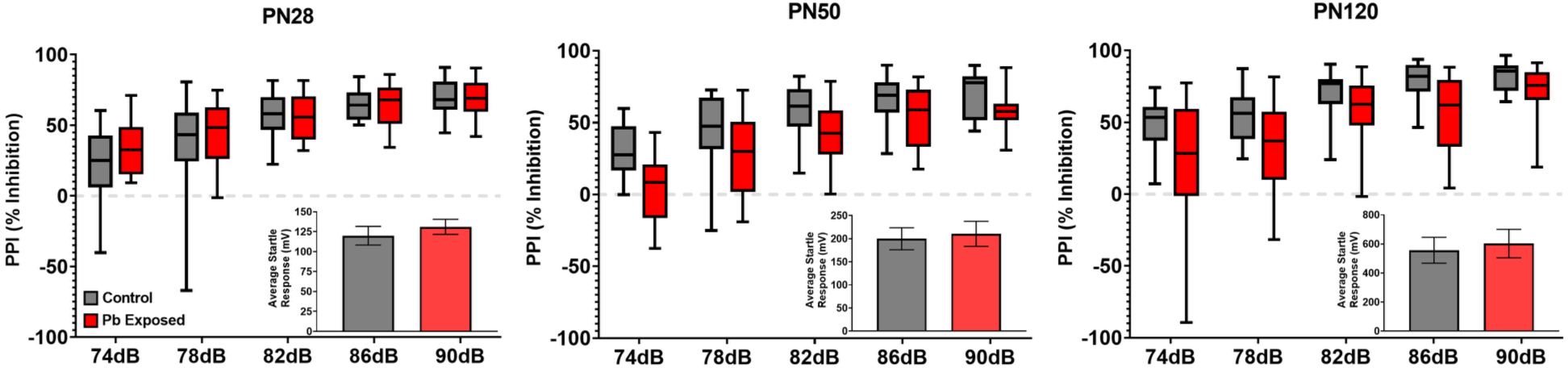
Prepulse inhibition was disrupted in adult male rats exposed to Pb^2+^. There was no effect of Pb^2+^ exposure at PN28, but there was marked inhibition of PPI in male rats at PN50 and PN120. Insert graphs in each panel show the startle response at 120 dB. No effect on startle response was observed in Pb^2+^-exposed male rats relative to controls at any age (p’s > 0.10). Thus, CELLE does not alter the animal’s sensitivity to a startle. However, CELLE presented with significant deficits in PPI in male rats relative to controls at PN50 and PN120 (p’s < 0.01).

## Discussion

### Summary of Main Findings

Childhood Pb^2+^ intoxication alters the neurodevelopmental trajectory of the brain, including the HC and PFC, and increases the risk of neurodevelopmental disorders, such as schizophrenia (see Albores-Garcia et al., 2021) and epilepsy (Sasmaz et al., 2003). Here, we asked how Pb^2+^ exposure impacts the HC at the network level using a rodent model of chronic early-life Pb^2+^ exposure (CELLE) that presents with molecular and cellular correspondences to schizophrenia (Stansfield et al., Transl Psych 5: e522, 2015). The prediction, based on established neurobiological deficits resultant of Pb^2+^ exposure, including hypofunction of the NMDA receptor (Guilarte, 1997), reduced LTP (Nihei et al., 2000), and reduced entorhinal cortex inputs (Zhou et al., 2023), is that chronic Pb^2+^ exposure should diminish HC activity. This outcome would help explain Pb^2+^ exposure-induced cognitive deficits observed in CELLE rats (McGlothan et al., 2008; Kuhlmann et al., 1997; Jaako-Movits et al., 2005) and potentially the cognitive and intellectual deficits seen in children. Simply put, reduced HC activity would reduce HC function.

Alternatively, the loss of PVGI interneurons (Stansfield et al., 2015) and impaired LTD (Zhao et al., 1999) predict network overexcitation in the HC. Instead of either prediction, we found that Pb^2+^ exposure caused network *hypersynchrony,* which appears as changes in the coordinated rhythms that enable efficient network processing. This suggests that Pb^2+^ exposure causes a rhythmic architectural shift in the HC network, reshaping activity patterns in a way that diminishes function. Specifically, we found that Pb^2+^ exposure resulted in four main HC network changes, including: (1) HC theta hypersynchrony, (2) gamma oscillation amplification, (3) disruption of behavioral-related theta and gamma network modulations, and (4) a striking exacerbation of absence seizures characterized by maximal theta synchrony during SWDs. We also showed that HC SWDs corresponded to SWDs occurring in cortical mPFC sites. Lastly, we predicted that aberrant HC hypersynchrony and SWDs caused by CELLE would impair sensorimotor gating because it is disrupted by aberrant HC activity (Zhang et al., 2002). Thus, we tested Pb^2+^-exposed rats on the PPI of the acoustic startle reflex task. We found that PPI was disrupted in late adolescence and young adult male rats but not female or juvenile males—a pattern that recapitulates PPI deficits in schizophrenia (Kumari et al., 2004). Theoretically, these results suggest that HC hypersynchrony and SWDs cause behavioral deficits by reducing the flexibility and processing efficiency of the HC network and causing behavioral arrests (a.k.a., the blank stare) via intermittent absence seizures (Holmes et al., 1987; Taylor et al., 2019). That is, hypersynchronous HC networks, the extreme of which is seizure-related activity, could be causing some of the Pb^2+^ exposure-related PPI and cognitive deficits in CELLE rats and possibly in Pb^2+^ exposed children (Kponee-Shovein et al., 2020).

### How can hypersynchrony cause network dysfunction?

Oscillations coordinate HC neuronal ensembles, creating time windows for synaptic integration and encoding dimensions of ongoing, remembered, and imagined future experiences, in turn guiding spatial navigation and decision-making processes (Buzsaki & Watson, 2012; Buzsaki & Wang, 2012). In addition, HC encoding of the present location and possible future trajectories comprise sequential neuronal activations that reiterate and update gamma-linked spatial representations, which occur on every cycle of the theta rhythm (Lowet et al., 2023; Buzsaki & Wang, 2012). Whereas gamma-theta synchrony usually enables cognitive representations, any change (in any direction) in the spatiotemporal structure of these neural representations and rhythmic mechanisms would diminish neural computations and mnemonic processes. Our finding of theta-gamma hypersynchrony in Pb^2+^ exposed rats argues that ‘overly’ structured neural activity may diminish HC function, presumably by reducing the variability and flexibility of spiking activity patterns to a narrower range of less dynamic network states. For example, information processing is lost if all neurons are synchronous (or asynchronous). This suggests there must be an optimal point of rhythmic coordination to maximize processing, which is assumed to be benchmarked by the control rats. In support of this notion, several neuropsychiatric disorders are associated with hypersynchrony, including epilepsy (Katuri et al., 2023), schizophrenia (Delgado-Sallant et al., 2021), Alzheimer’s disease (Kazim et al., 2021), among others.

### Theta and SWDs

HC SWDs reflect entrainment to a corticothalamic circuit oscillation initiated in the cortex and amplified by intrathalamic circuits. The robust theta-band fundamental frequency of SWDs suggests the involvement of normal theta rhythmogenic mechanisms, including but not limited to interneuron generators and modulators (Amilhon et al., 2015), medial septal GABAergic input (Borhegyi et al., 2004), and Huygens synchronization (Kocsis et al., 2021). We also found that theta frequency was lower in Pb^2+^-exposed rats during running and immobility. Slower frequency oscillations are generally associated with synchronization of larger populations of neurons. Lower frequency theta could, therefore, reflect the broader synchronization of HC networks, entraining a higher proportion of neurons and restricting HC functional capacity by reducing the number and flexibility of network states available. Here, the reduction in GABAergic diversity (Stansfield et al., 2015) (loss of PVGI cells) in chronic Pb^2+^ exposure, specifically within the context of homeostatic regulation of the E/I balance, suggests that the theta-gamma hypersynchrony observed may be due to an overrepresentation of the contribution of one inhibitory interneuronal subtype to the network dynamics recruiting larger populations of neurons. It is provocative to consider that the theta-band fundamental frequency of SWDs may reflect the pathological engagement of theta rhythmogenic mechanisms that are normally the foundation of spatial encoding and episodic memory systems. We argue that disrupted theta rhythmic coordination of HC networks is a key mechanism of chronic Pb^2+^ exposure-related cognitive deficits.

### Possible translational avenues to explore

Given our results, electrophysiological measurements (e.g., EEG, MEG, deep electrodes) should be helpful in distinguishing endophenotypes in Pb^2+^ exposed populations, as hypersynchrony and SWDs should be reflected, in some way, in these approaches (e.g., Kmiecik-Malecka et al., 2009). This strategy enables mappings of clinical features onto underlying disease mechanisms based on the generalization that similar clinical presentations are likely to share similar underlying pathophysiology. When distinct clusters among features can be identified, it serves the purpose of identifying a mechanism of particular importance, because the manifestation of the disease depends on it, or divides the disease into subtypes. The significance of this for distinguishing endophenotypes is that the development of effective treatments of the complex cognitive effects of Pb^2+^ exposure may also exhibit efficacy for cases of other diseases that share a hypersynchronous profile stemming from disruption of GABAergic dynamics.

Theta hypersynchrony and SWDs suggest possible interventions for hypersynchronous neural activity resulting from Pb^2+^ exposure, including (1) deep brain stimulation in a network hub to disrupt hypersynchrony and circuit entrainment to rhythmic activity (Zangiabadi et al., 2019), (2) pharmacological interventions to reduce excitability and rhythmogenesis (e.g., anticonvulsants), or (3) using the promising BDNF-mimetic 7,8-dihydroxyflavone (7,8-DHF) which mitigates the negative impact of Pb^2+-^exposure on presynaptic terminals (Zhang et al., 2018). This suggests that 7,8-DHF might modify network dynamics with a safe, inexpensive substance found in many fruits in the human diet (Guilarte, 2023). All these approaches require further research on the efficacy and optimal timing of each intervention to reduce hyperexcitability and enhance cognitive function in cases of chronic Pb^2+^ exposure, as well as their long-term impacts on brain function and cognitive/behavioral health.

### Conclusion

Chronic Pb^2+^ exposure during early brain development leads to significant alterations in HC network activity, including theta-gamma hypersynchrony and absence seizures. Theoretically, aberrant synchrony disrupts informational processing in the HC, resulting in diminished cognitive functions and impaired sensorimotor gating. Interestingly, these deficits may manifest more subtly in children as blank stares, distractibility, and reduced intellectual abilities in the classroom setting. If left untreated, such issues may lead to disruptive behaviors in the short term and social delinquency in the long term.

## Acknowledgements

This work was supported by R01 ES006189 and R01 ES035827 to T.R.G. and R01 MH113626 to T.A.A. We thank the members of the Guilarte Lab and the Allen Lab for their assistance with experiments and for providing constructive feedback on the manuscript. Acknowledging AI: ChatGPT-4-turbo (OpenAI) was used sparingly to help with snippets of MATLAB scripts, SciSpace was used to help with literature searches, and Grammarly was used to edit text for punctuation and grammar. AI was not used to generate content.

---

**Supplemental Figure 1.**
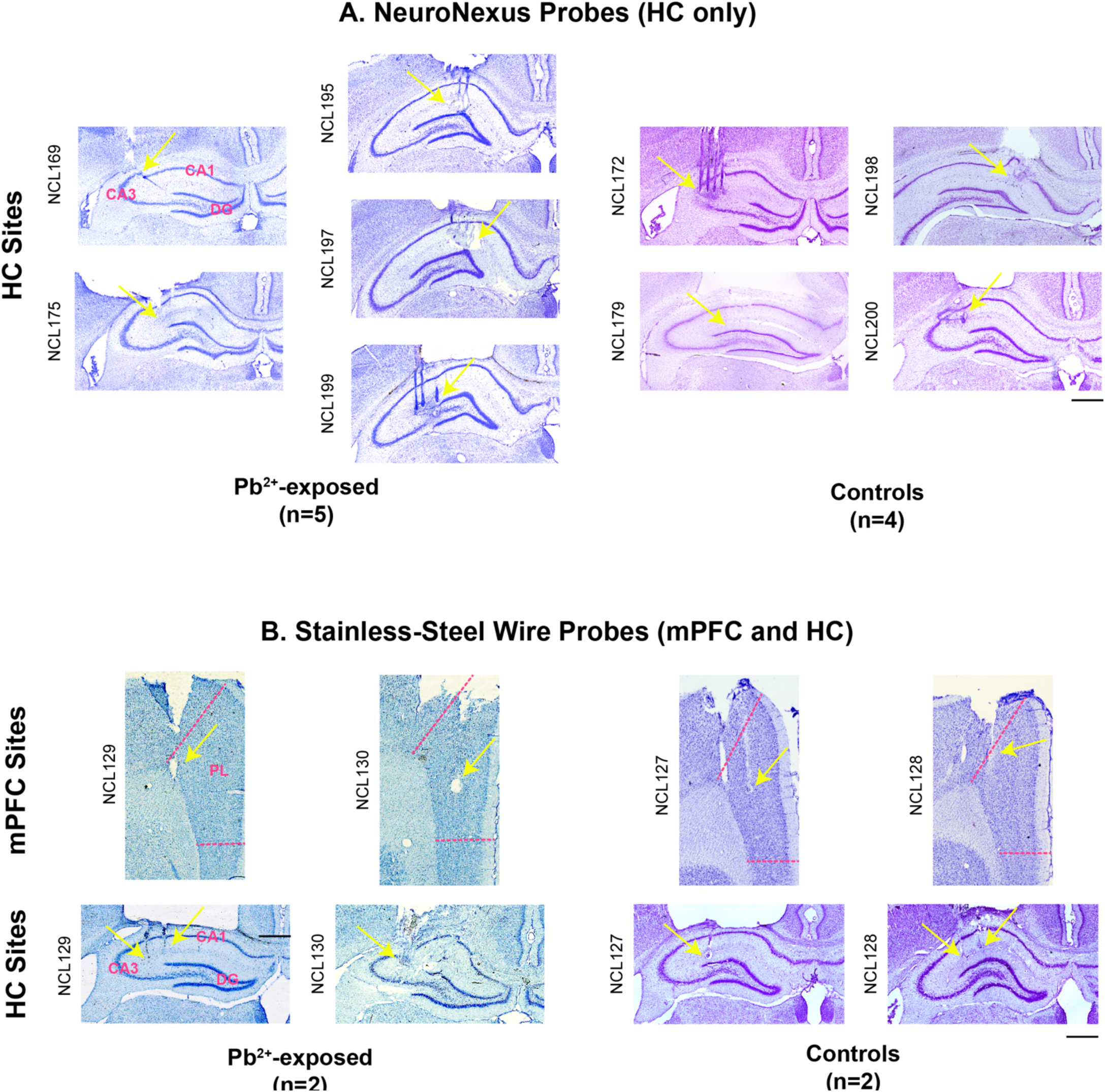
Histological estimation of recording sites. Nissl-stained coronal brain sections are presented for all rats. The yellow arrows indicate electrode markings in each section. **A.** In nine rats, NeuroNexus probes (4 shanks) were implanted in the HC. Pb^2+^ exposed rats are in the upper left cluster, while controls are in the upper right cluster. **B.** In four rats, custom-made stainless-steel (50μm tips) dual-site electrodes were implanted in the mPFC and HC. Pb^2+^ exposed rats are in the lower left cluster, while controls are in the lower right cluster. The cal bars are 1mm. Abbreviations: CA1, field CA1 of the HC; CA3, field CA3 of the HC; DG, dentate gyrus of the HC; and PL, prelimbic region of the mPFC.

**Supplemental Figure 2.**
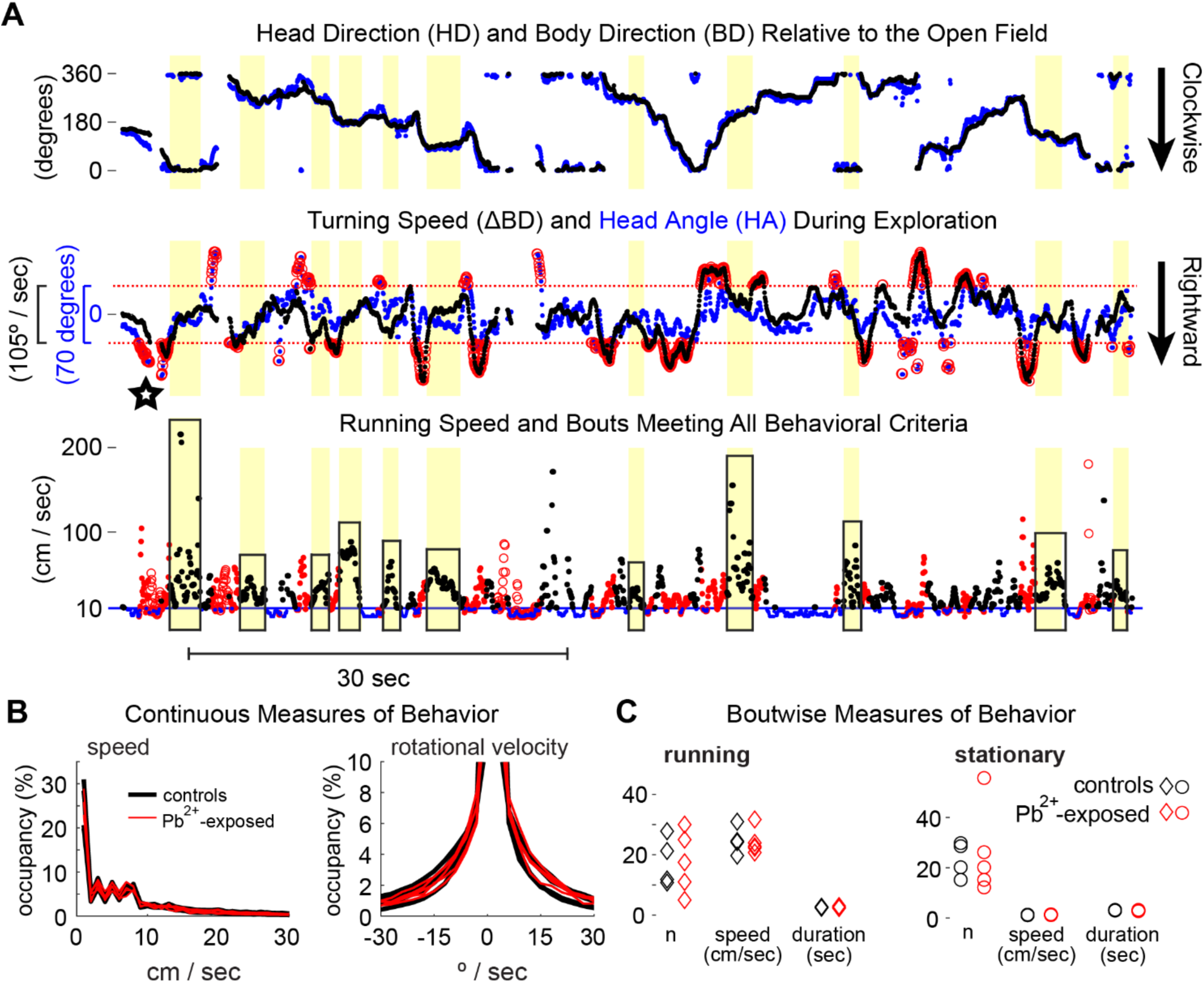
Precise delineation of stationary and running bouts. **A.** Head and body direction (**top**) were used to derive rotational velocity and head angle (**middle**), which in turn were used to identify bouts of running (boxed at **bottom**) while excluding rotational and reorienting behaviors (red points and circles). **B.** Speed occupancy (**left**) and rotational velocity (**right**) measured throughout behavioral sessions did not differ between control and Pb^2+^-exposed rats, nor did bout-wise behavioral measures (**C**) of running (**left**) and stationary bouts (**right**), including the numbers of bouts, median movement speeds, and bout durations.

**Supplemental Figure 3.**
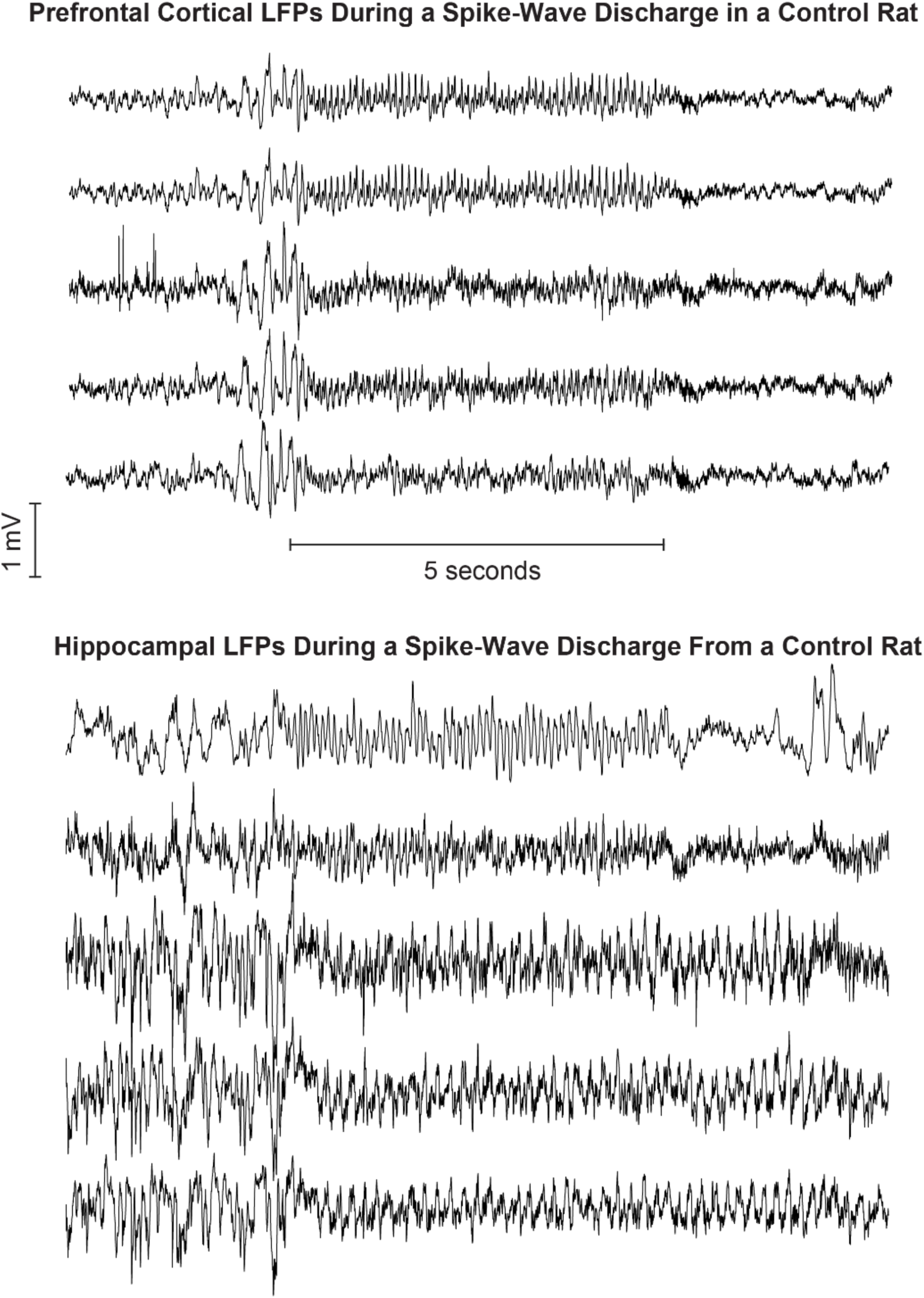
SWDs in a control rat. An example of an SWD recorded simultaneously in mPFC (above) and HC sites (below). The HC sites showed some variability in the LFP waveforms during the seizure and the continuation of theta oscillations following seizure offset. mPFC SWDs only showed up when HC SWDs did, not vice versa.

**Supplemental Figure 4.**
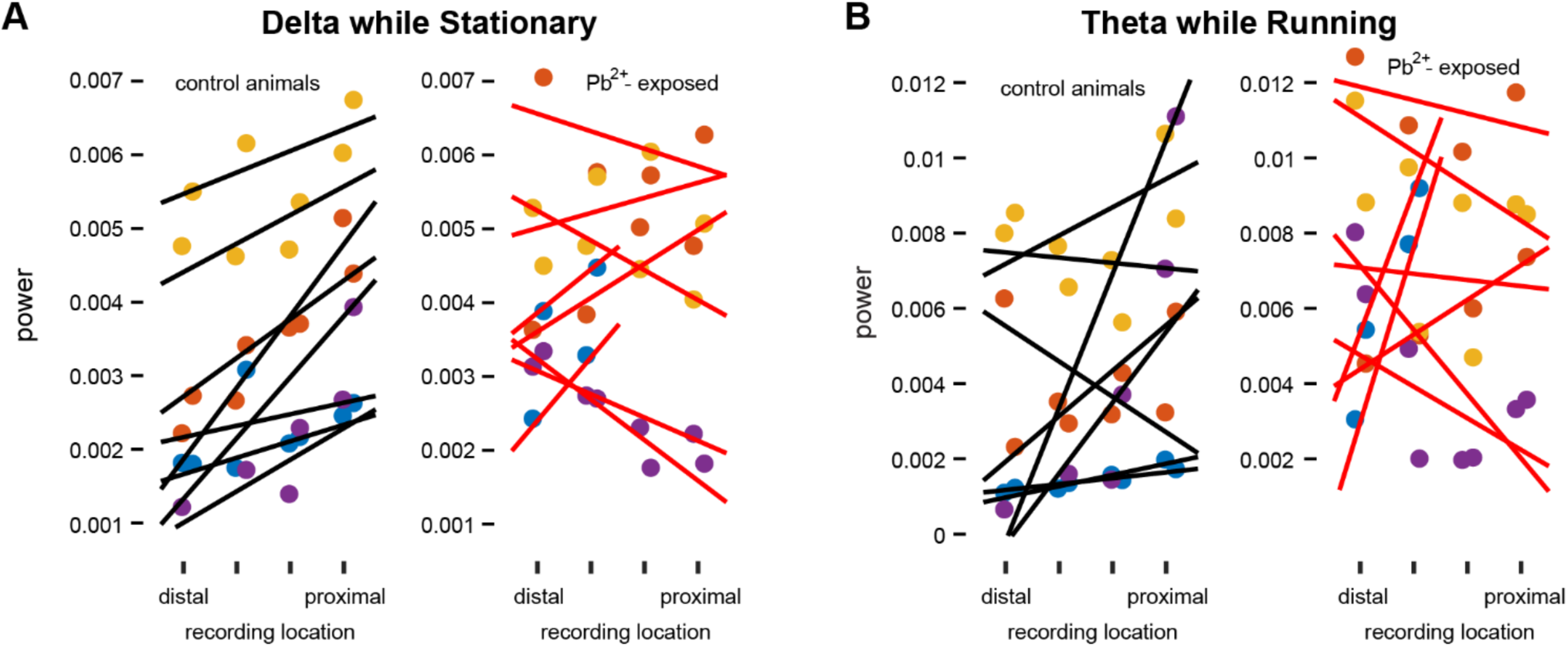
Proximal-distal gradient of delta and theta power in HC. **A.** An apparent proximal-distal gradient in delta power across probe shanks in stationary control rats (**left**), was absent in Pb^2+^-exposed rats (**right**). Black and red lines in A and B reflect the best fit to power estimates along each rat’s dorsal and ventral rows of tetrodes (across shanks). **B.** Theta power at sites along the proximal-distal axis in control and Pb^2+^-exposed rats during running bouts.

**Supplemental Figure 5.**
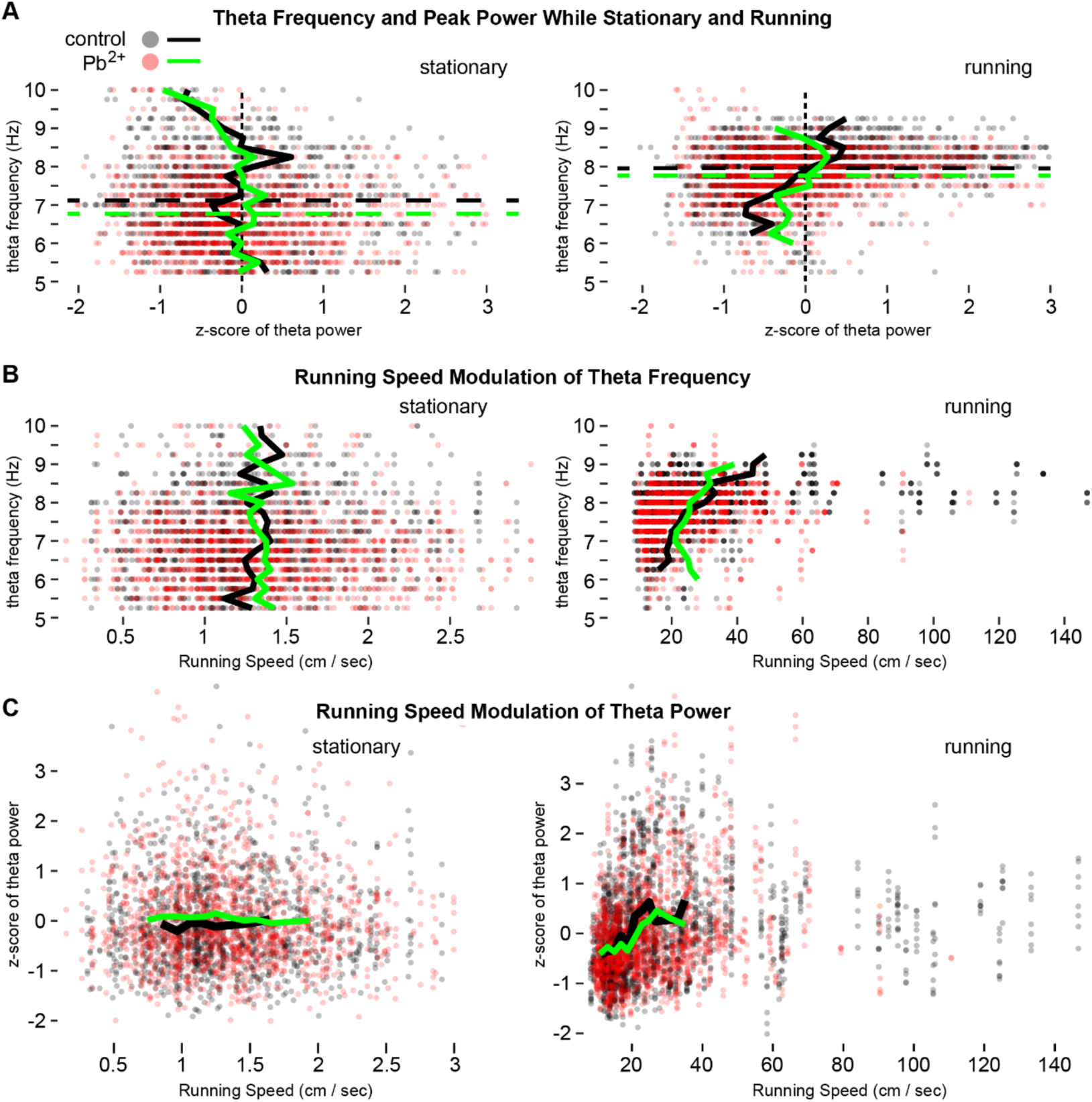
Pb2^+^ exposure reduces theta frequency and disrupts running speed modulation. **A.** Scatter plots of theta power and frequency taken from the power spectral density for each recorded channel during each bout of stationary (**left**) and running behaviors (**right**) for all rats. Black and green lines indicate means for control and Pb^2+^-exposed rats, respectively. Dashed lines indicate the mean theta frequencies for each group. **B.** Running speed modulation of theta frequency. **C.** Running speed modulation of theta power.

**Supplemental Figure 6.**
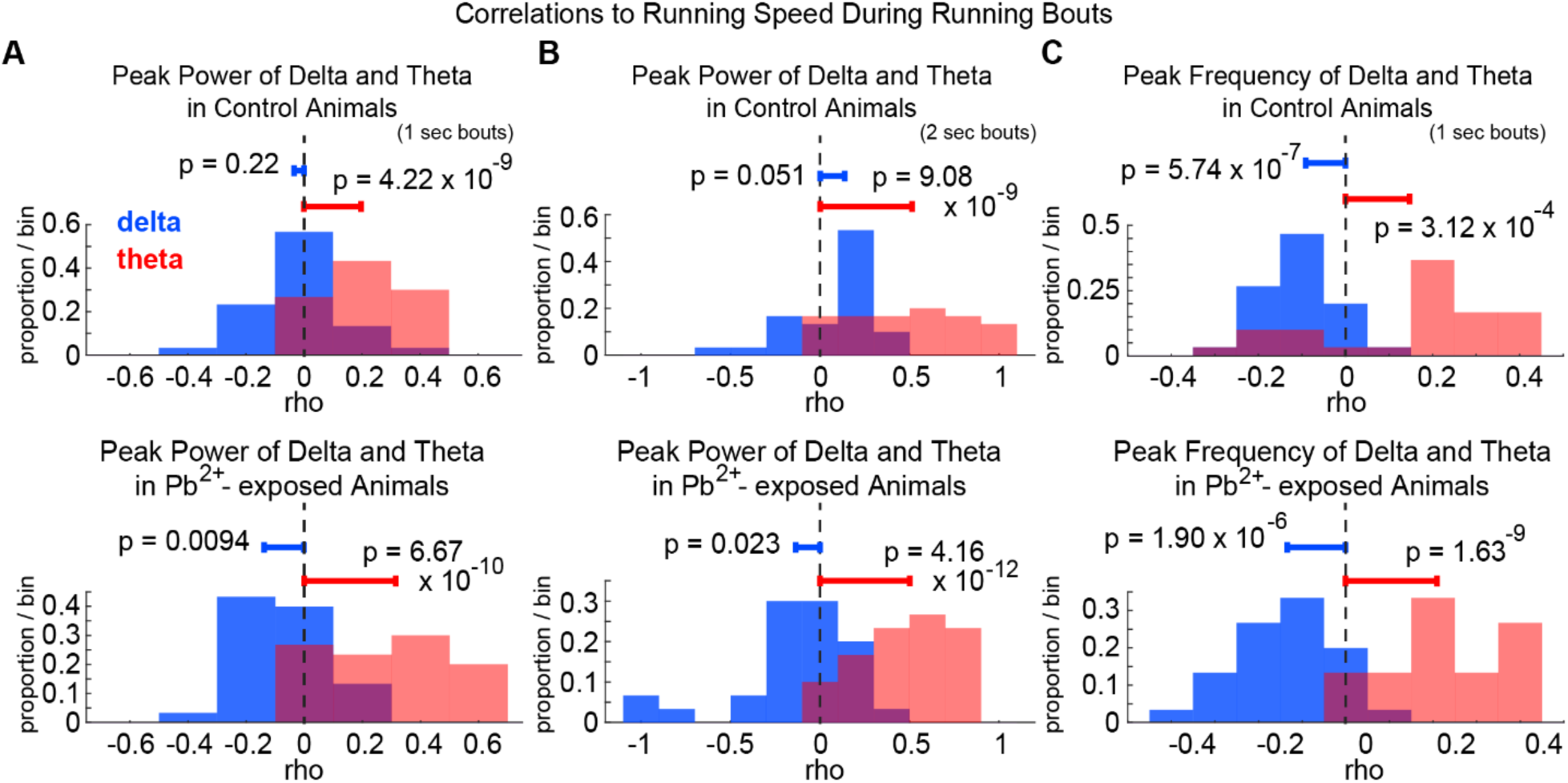
Running speed modulation of theta power is elevated in Pb^2+^-exposed rats. **A.** Distributions of correlation coefficients between delta (blue) and theta (red) power estimates and running speed (one-sample t-tests, null = 0). Individual correlations were calculated across running bouts (1 sec) for each recorded channel from control rats (**top**) and Pb^2+^-exposed rats (**bottom**). **B.** As in A, for 2-sec running bouts. **C.** As in A, but comparing the delta and theta peak frequencies correlations to running speed. Significance as indicated.

**Supplemental Figure 7.**
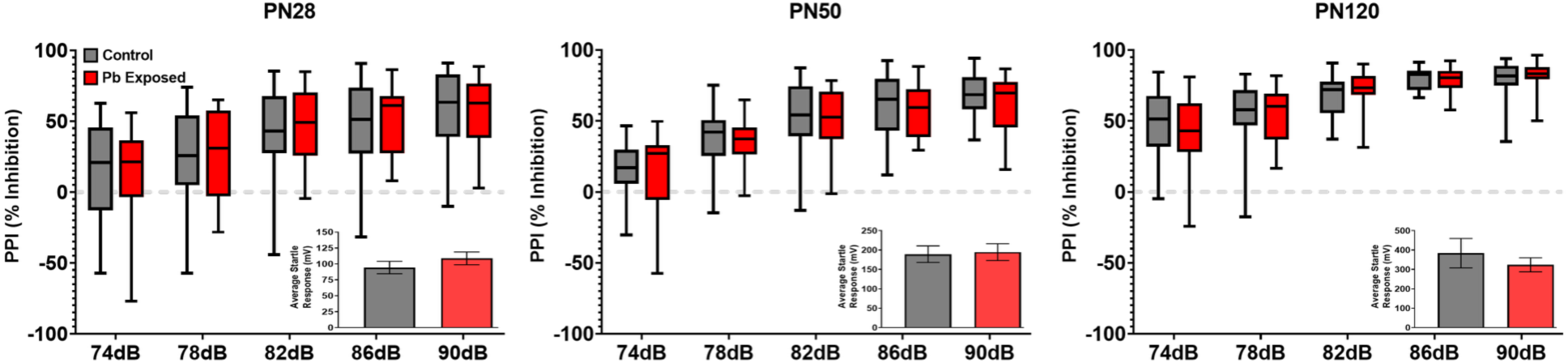
Prepulse inhibition is not disrupted in adult female rats exposed to Pb^2+^. No effect of Pb^2+^ exposure was found in female rats at any age (p’s> 0.10). Insert graphs in each panel show the startle response at 120 dB. No effect on startle response was observed in Pb^2+^-exposed female rats relative to controls at any age (p’s>0.10), nor was there an effect of Pb^2+^ relative to controls (p’s > 0.10).

**Supplemental Table 1.** Blood Pb^2+^ levels (µg/dL) in male and female rats contributed to the data in Fig5 and SupplFig7, respectively. Each value is the mean ± sem. Sample size n’s = number of litters (one male and female animal per litter).

| AGE | PN28 |  | PN50 |  | PN120 |  |
| --- | --- | --- | --- | --- | --- | --- |
| SEX | Males | Females | Males | Females | Males | Females |
| Control | ≤1.9<br>n=18 | ≤1.9<br>n=16 | ≤1.9<br>n=17 | ≤1.9<br>n=15 | ≤1.9<br>n=17 | ≤1.9<br>n=15 |
| 1500ppm<br>PbAc | 21.1 ± 0.9<br>n=18 | 20.9 ± 1.1<br>n=16 | 20.2 ± 1.1<br>n=17 | 22.1 ± 1.3<br>n=20 | 19.6 ± 1.3<br>n=17 | 24.3 ± 2.2<br>n=14 |
| p-value | ≤0.0001 | ≤0.0001 | ≤0.0001 | ≤0.0001 | ≤0.0001 | ≤0.0001 |

